# GPRC5A regulates keratinocyte adhesion and migration through nuclear translocation of its C-terminus region

**DOI:** 10.1101/2023.11.28.569012

**Authors:** Sarah Chanteloube, Choua Ya, Gabrielle Le Provost, Aurore Berthier, Cindy Dieryckx, Sandrine Vadon-Le Goff, Florence Nadal, Bérengère Fromy, Romain Debret

## Abstract

G-Protein Coupled Receptor, Class C, Group 5, Member A (GPRC5A) is well-documented in lung and various epithelial cancers. However, its role in the skin remains unexplored. In this study, we investigated the function of this receptor in skin biology and our research demonstrated that its expression responds to mechanical substrate changes in human primary keratinocytes. Furthermore, we observed GPRC5A reinduction during wound healing at the leading edges in an *ex vivo* burn model, coinciding with the translocation of its C-terminal region into the nucleus. We identified the cleavage site of GPRC5A by N-TAILS analysis, and cathepsin G was characterized as responsible for proteolysis in cultured cells.

To gain a deeper understanding of GPRC5A’s role in keratinocyte, we performed GPRC5A knockdown in N/TERT-1 cells using short-hairpin RNA. Our findings strongly suggest a close association between GPRC5A and adhesion regulation pathways, but also demonstrate that GPRC5A^KD^ enhanced cell adhesion while reducing cell migration and differentiation. Importantly, these effects were reversed by adding a recombinant polypeptide mimicking the C-terminal region of GPRC5A.

Overall, our study reveals an unexpected role of GPRC5A in regulating keratinocyte behavior, implicating its C-terminal region translocation into the nucleus. These results open up interesting strategic pathways for wound healing.

## INTRODUCTION

In addition to the biomechanical properties of extracellular matrix (ECM), mechanical cues influence cell behavior. Mechanotransduction corresponds to the conversion of mechanical information to biochemical cascades and has been involved in regulating keratinocyte functions to maintain epidermal homeostasis as it can regulate their proliferation, differentiation, morphology, and migration (Jaalouk & Lammerding, 2009; Wang *et al*, 2009; Wong *et al*, 2011). *In vitro* studies showed that keratinocytes respond to mechanical signals and may dictate how skin and cutaneous wounds interact with the environment (Lumpkin & Caterina, 2007) (Mikesell *et al*, 2022). Substrate stiffness was demonstrated to promote keratinocyte proliferation and migration, whereas it inhibits the differentiation process (Wang *et al*, 2012). Moreover, the substrate stiffness also regulates the epidermal barrier through Jun kinase pathway phosphorylation which contributes to adherent junction formation (You *et al*, 2013). According to these observations, keratinocyte mechanotransduction is a crucial process in epidermal homeostasis and an improper process can cause many pathological changes, including impaired wound healing (Jaalouk & Lammerding, 2009). Chen et al. reported the relationship between stiffness and skin wound healing and demonstrated that optimal stiffness is necessary to improve wound healing (Chen *et al*, 2016). The ECM stiffness gives favorable conditions and support for the initiation of keratinocyte migration, whereas abnormal ECM mechanical properties lead to defects in skin repair (Kenny & Connelly, 2015). Immediately after the injury, the basement membrane disappears, and the keratinocytes start to migrate on the underlying dermis ECM mostly composed of blood platelets and fibrine. This clot will be rapidly replaced by the fibroblasts in an ECM rich in fibronectin, tenascin C and type III collagen to form the granulation tissue (Diller & Tabor, 2022). Hinz et al. showed on rat wounded skin that the provisional matrix increased from 10kPa in the healthy dermis to 18 kPa at 7 days after wounding and 50 kPa at 12 days, when measuring the fibrotic tissues and contracting wound granulation tissue using AFM (atomic force microscopy) device (Hinz, 2010).

More particularly in the epidermis, the mechanotransduction system includes the participation of several protein complexes such as the cytoskeletal elements (e.g. F-actin, intermediate filament, and microtubules), mechanically activated ion channels (e.g. Connexin, Piezo, calcium sensing receptors, transient receptor potential), focal adhesions, integrins, adhesion receptors, and G protein-coupled receptors (GPCRs) (Wong *et al*., 2011). Several studies have indicated that some GPCRs can be activated by mechanical forces, such as cellular stretching or compression in endothelial cells and cardiac tissue (Erdogmus *et al*, 2019; Lin *et al*, 2022; Xu *et al*, 2018; You *et al*., 2013). Investigations conducted on keratinocytes have provided evidence that GPCRs regulate cell proliferation, differentiation, and migration (Aragona *et al*, 2017; Liu *et al*, 2014; Pilar Pedro *et al*, 2023). However, it is noteworthy that, despite their apparent roles in epidermal homeostasis, no current study has conclusively demonstrated a direct response of GPCRs to mechanical forces within keratinocytes.

In the present study, we provide evidence that a specific GPCR, GPRC5A, plays a pivotal role in governing human keratinocyte behavior by modulating both the adhesion and migration processes. Furthermore, GPRC5A seems intricately involved in epidermal differentiation, as its deficiency disrupts proper stratification. Our working hypothesis posits that keratinocytes exhibit responsiveness to alterations in tissue stiffness during wound healing, thereby triggering the initiation of the reepithelization phase. Hence, we explore here, using *in vitro* and *ex vivo* human model, the role of GPRC5A in the early stages of wound healing to enhance the wound re-epithelialization process.

## RESULTS

### Searching for new mechanosensitive candidates in human primary keratinocytes (2D)

To identify new mechanosensing receptors involved in keratinocyte mechanotransduction, we performed a genome-wide expression analysis by RNA-sequencing of human primary keratinocytes cultured on glass surface (Glass), and on a soft substrate (Soft) by using polyacrylamide hydrogels. All cells were culture-conditioned for 3 days and cells on soft hydrogels were also re-plated on glass surface (Soft_glass) for 24 h (**Figure 1**). Analysis revealed a strong signature of Glass condition compared to Soft and Soft_glass conditions (**Figure 1a**). However, 43 genes were specifically overexpressed when cells where replated on Glass (**Figure 1b**). One of the most upregulated gene was the G protein-coupled receptor class C family 5 member A (GPRC5A) (**Figure 1c**), suggesting a candidate responsive for keratinocyte mechanotransduction. To confirm this, GPRC5A expression was analyzed after a 72 h preconditioning on different substrate rigidities (soft, medium, and rigid hydrogels or glass), prior re-plating on glass substrate for 24 h. GPRC5A expression was significantly overexpressed during the transition from hydrogels to glass substrate while no significant difference was observed from glass to glass replating (**Figure 2d-g**). Consequently, the more the difference between initial and final substrates, the stronger the expression of GPRC5A (up to 6 times increase between the soft and the glass conditions). Taken together, these first results reveal a mechanosensitive role of the GPRC5A receptor in human keratinocyte.

**Figure 1:**
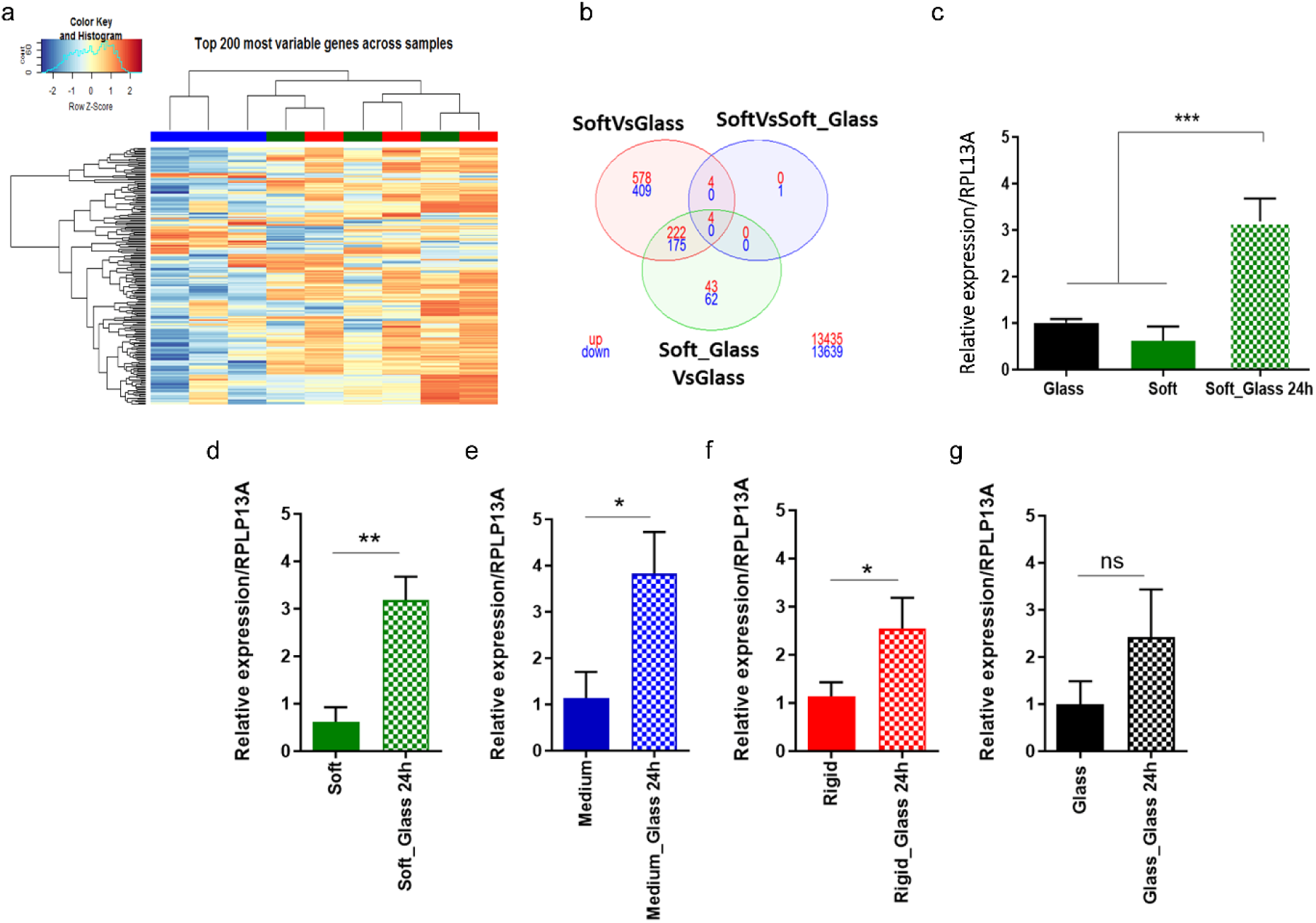
Transcriptomic analysis: GPRC5A expression is correlated to stiffness changes. (a) Heatmap of differentially expressed genes (FDR < 0.05) by comparing Glass (blue), soft (green) and soft_glass (red) conditions. (b) Overlap between differentially expressed genes (using gene symbol) from the analysis shown in (a), displayed as Venn diagram. (c) qRT-PCR validation of one of the topmost significant gene showed in (b): GPRC5A. One-way ANOVA, n=3 ; *** P< 0.001. (d-g) mRNA expression of GPRC5A on soft (d), medium (e), rigid stiffness (f), and glass surface (g) for 3 days and after re-plating on glass substrate. T-Test; * P< 0.05. Results are presents as means ± SD and were performed in at least three independents biological replicates.

**Figure 2:**
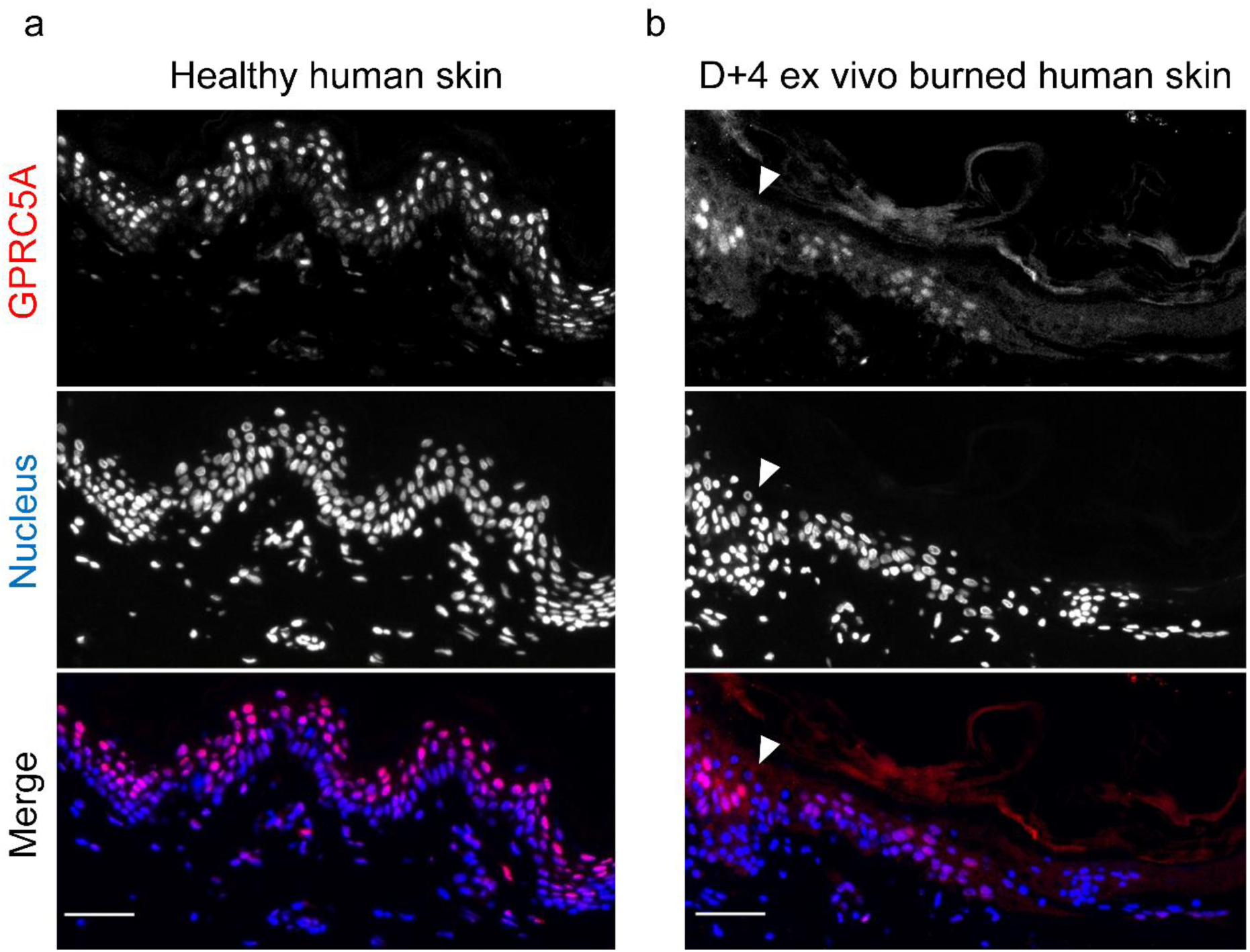
Effect of healing on cutaneous GPRC5A expression. GPRC5A immunostaining (red) in human healthy abdominal skin (a) or after 4 days post-burn (150°C, 3 seconds) in *ex vivo* abdominal skin (b). Nucleus were counterstained with DAPI (blue). The white arrow stands for the edge of the burn wound. Scale bars: 50 µm.

### GPRC5A expression in homeostatic and altered human skin

GPRC5A is predominantly localized peri- and intra-nuclearly in the differentiated healthy human epidermis (**Figure 2a**). However, observations 4 days after burn in an *ex vivo* human skin model have indicated that the receptor’s expression is transferred from differentiated layers to the basal migrating keratinocytes (**Figure 2b**), in a similar manner as previously observed by Aragona et al (Aragona *et al*., 2017), suggesting a potential role in the re-repithelialization process. Some spatiotemporal expression pattern was also noted: receptor expression was completely lost during the initial day post-burn and then restored after 4 days in culture, in migrating keratinocytes (**Figure S1**). Similar to the expression of GPRC5A, Keratin 14 disappeared entirely in the damaged area at D0. Subsequently, from day 4 to day 12, its expression was restored in the basal keratinocytes at the wound edges, corresponding to proliferating and migrating cells (**Figure S1**).

At this point, a noteworthy observation emerged when using two different antibodies to label each end of the GPRC5A receptor. While only cell cytoplasm was labeled by an anti-N-terminal GPRC5A antibody (anti-RAI3), an anti-C-terminal GPRC5A antibody (anti-GPRC5A) exhibited two distinct signals in the cytoplasm and in the nucleus respectively (**Figure S2**). Investigations using cultured primary human keratinocytes under different substrate conditions yielded similar results when transferring cells from soft to stiff substrates (**Figure 3a**).

**Figure 3:**
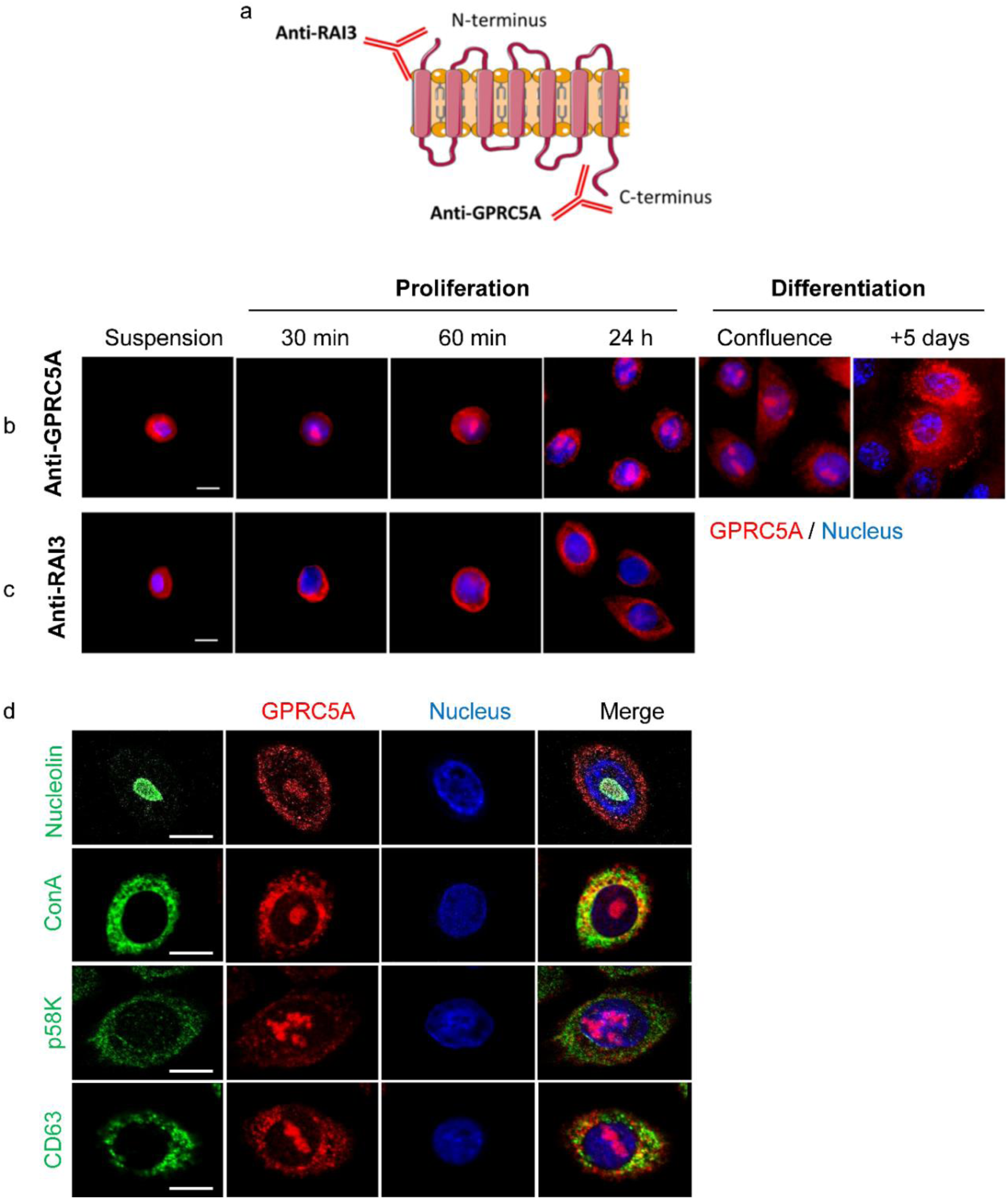
GPRC5A dynamic relocation during primary keratinocyte adhesion. (a) Graphic showing the antibodies recognition sites. (b-c) GPRC5A immunostaining (red) with anti-GPRC5A (b) and anti-RAI3 (c) antibody, and nucleus (blue). Scale bars: 10µm. (d) GPRC5A immunostaining (anti-GPRC5A, red), nucleus (blue) and the nucleolus (nucleolin), the endoplasmic reticulum (Con A), the Golgi apparatus (p58k) or the endosomal vesicles (CD63) in green. Scale bars: 25µm.

### GPRC5A translocation

To delve deeper into the role of GPRC5A in keratinocytes, the localization of GPRC5A was examined at various time points using two distinct antibodies (**Figure 3a**). We observed dynamic changes of GPRC5A during keratinocyte adhesion and differentiation. Notably, in cellular suspensions, both antibodies identified the receptor in a perinuclear compartment (**Figure 3b-c**). However, during the initial stages of cell adhesion and early differentiation, while the anti-N-terminal staining remained perinuclear, the anti-C-terminal antibody indicated a relocation of this region of the receptor into the nucleus. However, during the differentiation phase, GPRC5A C-terminal region expression within the nucleus decreased, seemingly returning to cytoplasmic vesicular compartments (**Figure 3b**). Subsequent co-immunolabeling experiments revealed co-localization of GPRC5A C-terminal region with specific endomembrane organelles such as the endoplasmic reticulum and the Golgi apparatus (**Figure 3d**).

This observation gains particular interest due to its concurrent timing with the upregulation of receptor expression following the transition from a soft substrate to a rigid one. Specifically, the receptor’s transcriptional increase extends up to 24 h after cells adhere to their substrate, while the translocation of GPRC5A the C-terminal region into the nucleus also begins within the first 30 minutes post-adhesion and persists until cell differentiation.

To date, the cleavage site of the C-terminal portion of GPRC5A has never been reported. GPRC5A cleavage was then analyzed in NHEK, 24 h post-seeding, using a N-terminal amine isotopic labeling (TAILS). Mass spectrometry (LC-MS/MS) analyses revealed 3 cleavage sites at positions 269, 321 and 336, with a more abundant cleavage site corresponding to the YAPY↓321STHF sequence (**Figure 4a**). Subsequent bioinformatic analysis identified the cathepsin G as a proteolysis candidate. To investigate this, we treated a recombinant peptide corresponding to the cytoplasmic C-terminal domain of GPRC5A (C-ter peptide) designed at the laboratory, with purified cathepsin G. Gel analysis following digestion revealed that the peptide could indeed be cleaved by the cathepsin G in less than 2 h under physiological conditions (**Figure S3**).

**Figure 4:**
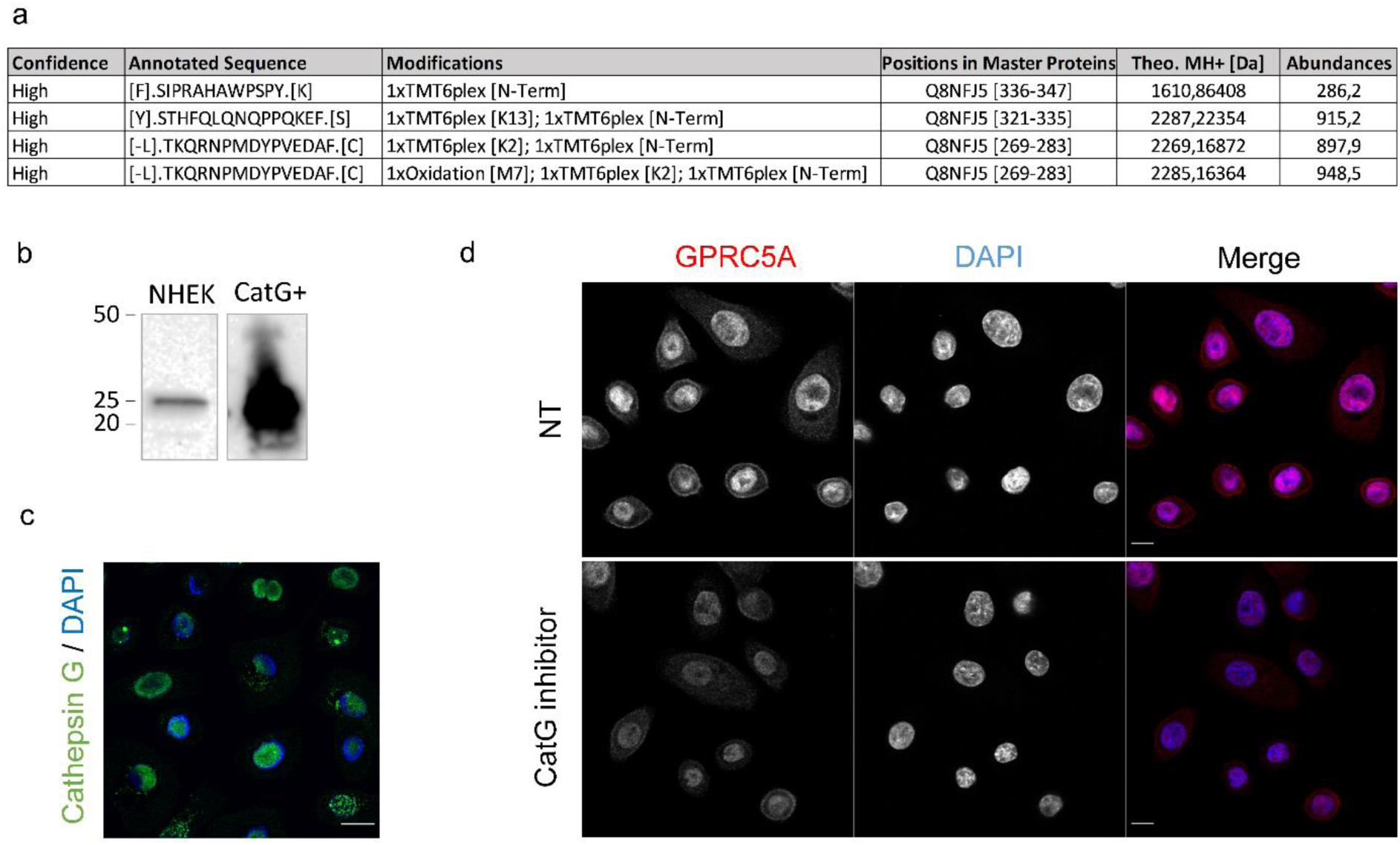
Analysis of the C-terminal cleavage of GPRC5A. (a) Results of MS analysis showing peptides corresponding to the GPRC5A receptor, 24 hours post-seeding in N/TERT-1 cells. [-L] corresponds to the end of the last intracellular domain of the endogenous GPRC5A. (b) Western-blot analysis of Cathepsin-G expression in primary keratinocytes (NHEK). Purified Cathepsin G (CatG+) is used as a positive control. (c) Cathepsin-G immunostaining (green) and nucleus (blue) in primary keratinocytes. Scale bar 20 µm. (d) GPRC5A immunostaining (red) and nucleus (blue) in primary keratinocytes +/- Cathepsin-G inhibitor treatment. Scale bars 10: µm.

To corroborate the presence of the cathepsin G protease, protein extracts from primary keratinocytes were subjected to western-blot analysis (**Figure 4b**). A 25 kDa band corresponding to purified cathepsin G was observed in keratinocyte extracts. This finding was further validated by immunofluorescence, where cathepsin G was detected in the perinuclear region of keratinocytes (**Figure 4c**).

To investigate the role of cathepsin G in the cleavage and translocation of the C-terminal part of GPRC5A, a specific protease inhibitor (shown as cell permeable and specific in dendritic cells (Reich *et al*, 2009)) was used on keratinocytes cultured on hydrogels with stiffness ranging from 4 to 14 kPa for 3 days before transfer onto glass slides to induce the nuclear translocation. Confocal microscopy confirmed nuclear translocation in non-treated cells, while the nuclear labelling was reduced after 4 h with the cathepsin G inhibitor (**Figure 4d**). This suggests the potential involvement of this protease in the cleavage of the C-terminal part of GPRC5A. Taken together, these results suggest that cathepsin G could be part in the cleavage of the C-terminal region of GPRC5A.

### GPRC5A impacts adhesion and cell survival pathways in keratinocyte

In order to understand the signaling pathways linked to GPRC5A in keratinocytes, a phosphoprotein array was performed following adhesion of GPRC5A knocked-down (KD) N/TERT-1 keratinocyte cell line (lentiviral infection of shRNA targeting GPRC5A sequence, sh-GPRC5A, or control sh-CTRL) 2 h and 24 h after replating on plastic surface. Repression of GPRC5A was validated at both RNA and protein levels (**Figure 5a**, and in primary keratinocyte using siRNA approach, **Figure S4**). The phosphorylation rate of 43 kinases and 2 related total proteins was analyzed (**Figure S5-6**). Histograms represent spot density of the most impacted kinases and the two total proteins (**Figure 6b-c**). This pointed out a two-fold increase in phosphorylation of JNK (jun N-terminal kinase), and a four-fold increase of FAK (Focal adhesion kinase) phosphorylation (Y397) in GPRC5A^KD^. A slight increase in phosphorylation of Src protein (Y419) was also observed, while β-catenin was not impacted by the lack of GPRC5A.

**Figure 5:**
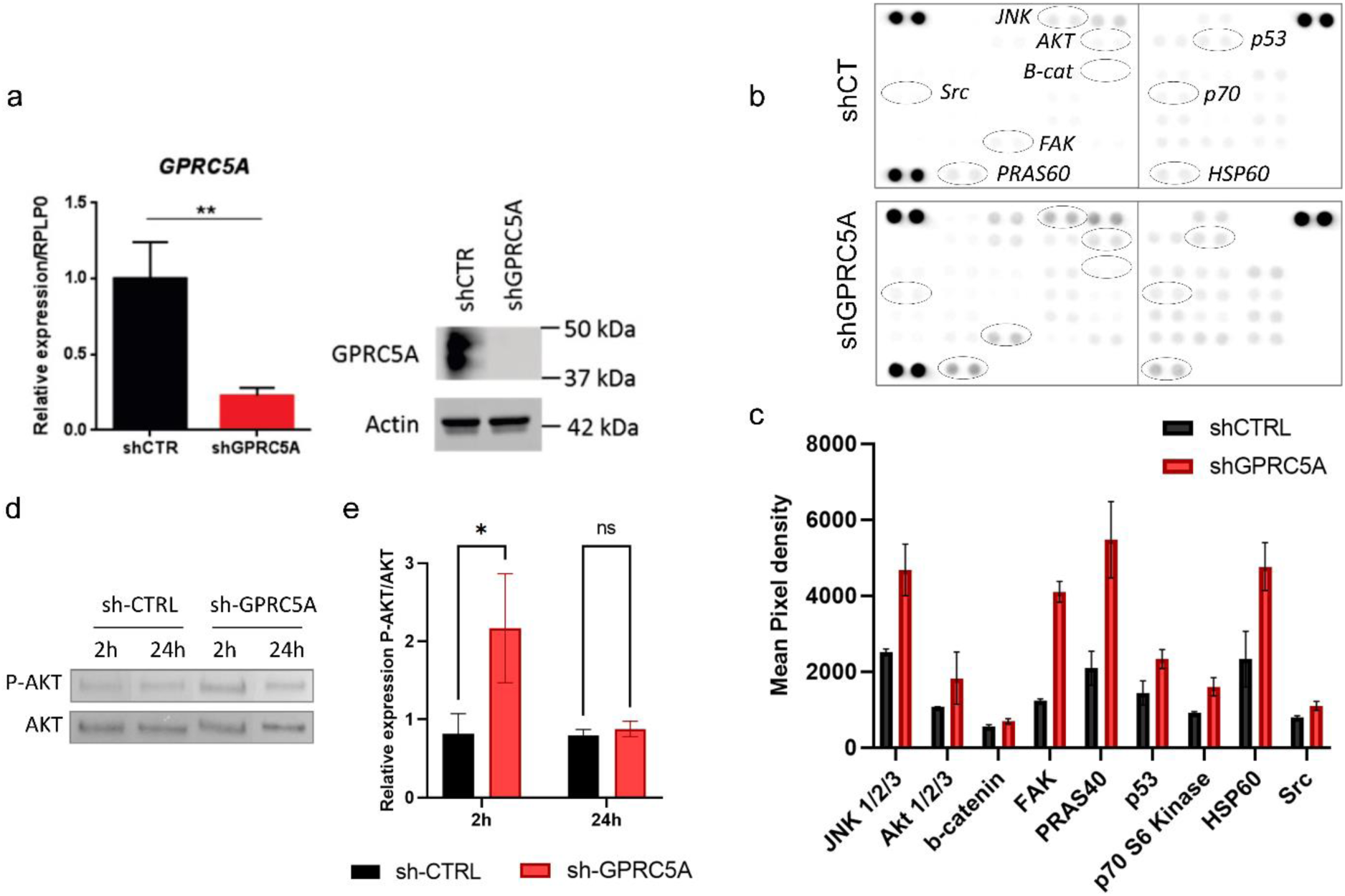
Protein phosphorylation level in N/TERT1 sh-GPRC5A (GPRC5A^KD^) vs sh-CTRL cells 24 hours post-adhesion. (a) Validation of GPRC5A inhibition performed through qPCR and Western-blot analysis. Tests were conducted on the stable immortalized human N/TERT-1 cell line using the GPRC5A-targeting shRNA (shGPRC5A) and a control shRNA (shCTR). One-way ANOVA test; T-test; ** P< 0.01. (b) Phosphorylation arrays 24 hours after adhesion. Each array was incubated with 200μg of cell lysate. Dark spots on array corners represent reference spots. (c) Chart showing the analysis of the most modulated kinase and two total proteins. (d) Western-blot analysis of Akt/P-Akt (S473, 60kDa) expression, in N/TERT1 sh-GPRC5A and sh-CTRL cells, 24 hours post-adhesion. (e) Quantification of the relative Akt phosphorylation over time, in sh-CTRL and sh-GPRC5A cells (n=2). Two-way ANOVA test; *p<0.05. All results are presents as means ± SD.

**Figure 6:**
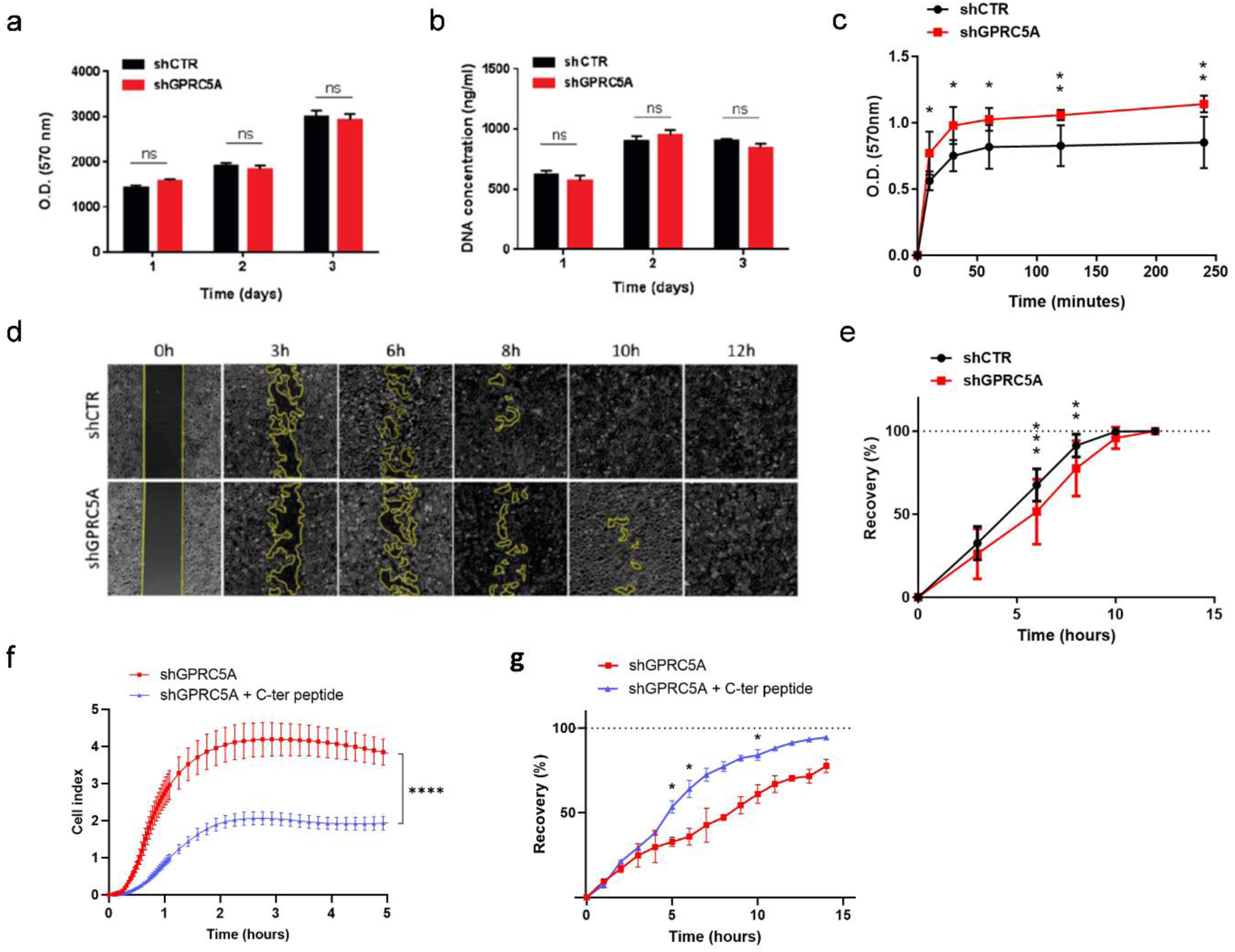
Knockdown of GPRC5A (GPRC5A^KD^) results in increased cell adhesion, but a reduced cell migration. (a-b) Proliferation assays in N/TERT-1 sh-GPRC5A and sh-CTRL cells: Alamar Blue (a) and DNA quantification (b). One-way ANOVA test, ns: non significative. (c) Quantification of adhesion assay (n=3) and (e) migration assay (n=3) in N/TERT-1 sh-GPRC5A and sh-CTRL cells. (d) Yellow line represents uncovered area during migration assay. Two-way ANOVA test; * P< 0.05, ** P<0.01 and *** P<0.001. (f-g) Mimetic C-ter peptide treatment effect (10 µg/ml, for 24 hours) on sh-GPRC5A cells. (f) Adhesion measured by impedance disturbing (n=4). Two-way ANOVA test; ****p<0.0001. (g) Quantification of migration assay (n=2). Two-tailed unpaired t-test; *p<0.05. All results are presents as means ± SD.

Additionally, we observed a three-fold increase of PRAS40 (T246) phosphorylation, and a two-fold increase of AKT (S473) and p53 (S15) phosphorylation in GPRC5A^KD^ cells. Finally, HSP60 total protein had a two-fold increase in GPRC5A^KD^ cells, which could be linked to a stressful environment for cells in the absence of the GPRC5A receptor. We noticed a time-dependent effect on protein phosphorylation, with an increase from the early steps of cell adhesion at 2 h post-adhesion (**Figure S6**), compared with the 24 h condition (**Figure 5c**).

To validate these findings, the experiment was replicated using new batches of cells, and Western blot analyses were conducted with a specific focus on AKT (**Figure 5d-e**). While the total protein concentration remained constant between the different samples, the labelling of S473 phosphorylation showed an increase in GPRC5A^KD^ cells (**Figure 5e**). As a result, these findings strongly suggest a close association between GPRC5A and signaling pathways regulating cell adhesion mechanisms.

### GPRC5A functionality in human keratinocytes

The study further investigated the effects of GPRC5A knockdown on keratinocyte behavior in vitro, focusing on proliferation, adhesion, and migration. To understand the role of the C-terminal domain in regulating these processes, a synthetic polypeptide mimicking this sequence of GPRC5A (C-ter peptide), which was effectively internalized by cells (**Figure S7**), was introduced to the knockdown cells.

To question whether GPRC5A knockdown could affect the keratinocyte proliferation process, we measured cell viability and DNA concentration over time. The results showed that GPRC5A inhibition does not affect cell proliferation (**Figure 6a-b** and **Figure S8**). According to our previous staining of GPRC5A at leading edges, we further tested whether GPRC5A deficiency could impair keratinocyte adhesion. We analyzed the ability of GPRC5A^KD^ cells and control cells to adhere to ECM components, such as type I collagen. Our results show that GPRC5A^KD^ cells adhered 2 times more efficiently than control cells, with a stronger effect 5 minutes after seeding that persists up to 120 minutes (**Figure 6c** and **Figure S8**).

Cell adhesion to the ECM is tightly linked with the cells’ ability to migrate during the wound healing process. We then analyzed the performance of GPRC5A^KD^ cells in a standardized migration assay. The results revealed that GPRC5A inhibition causes a reduced migration in comparison to the control (**Figure 6d-e** and **Figure S8**), proving causality between GPRC5A expression and adhesion/migration processes.

The role of the C-terminal part of GPRC5A was corroborated when GPRC5A^KD^ keratinocytes demonstrated a partial restoration of their adhesion and migration capacities after being treated with the recombinant peptide (**Figure 6f-g**). Indeed, both a decrease in keratinocytes adhesion, as well as an increase in their migration, were detected 24 hours after treatment.

These results demonstrate that GPRC5A is implicated in the regulation of keratinocyte adhesion and migration, but also that the C-terminal region of the receptor has a compensatory effect on these mechanisms, leading to faster wound closure during scratch assays.

To explore the function of GPRC5A in keratinocyte differentiation, we took advantage of 3D-reconstructed human epidermis (RHE), which provides a more physiological model of differentiation compared to monolayer cultures. GPRC5A expression was initially examined during the formation of reconstructed epidermis (**Figure 7a**). Immunostainings revealed that GRPC5A is expressed during the first week but decreases with longer times. This type of 3D differentiation model closely mimics both organogenesis and a form of wound healing, involving the proliferation and subsequent differentiation of keratinocytes. This transient expression pattern suggests a role for GPRC5A during the early stages of keratinocyte differentiation.

**Figure 7:**
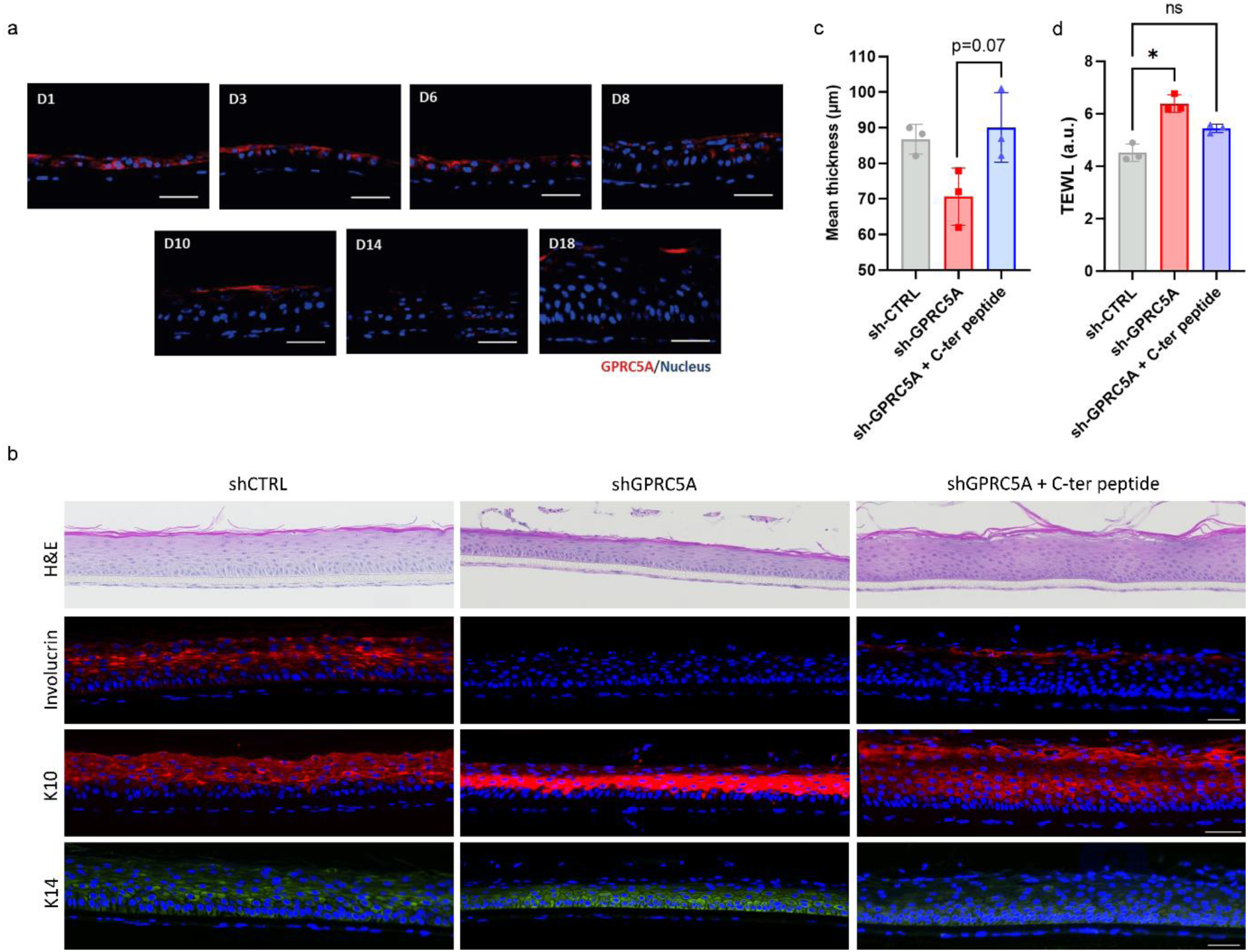
knock-out GPRC5A alters keratinocyte differentiation process. (a) GPRC5A immunostaining (red) in 3D-reconstructed human epidermis (RHE) at 1 to 18 days post-emersion. Nucleus was counterstained with DAPI (blue). Scale bars, 100 µm. (b) Haematoxylin-eosin staining (H&E) and immunofluorescence staining of involucrin (red), Keratin 10 (K10, red) and Keratin 14 (K14, green) in control and GPRC5A^KD^ RHE +/- treatment with GPRC5A C-ter peptide. Nuclei were counterstained with DAPI (blue). Scale bars, 50 µm. (c) Mean epidermis thickness (in µm), n=3. (d) RHE inside-out permeability using TEWL measurements (a.u: arbitrary unit), n=3. Kruskal-Wallis test, *p<0.05, ns= non-significant.

GPRC5A^KD^ cells were subsequently employed to generate RHE and evaluate the epidermal morphology after 16-days of air/liquid interface culture (**Figure 7b**). The results indicated that GPRC5A inhibition led to a decrease in epidermal thickness (**Figure 7c**). Immunostainings of keratinocyte differentiation markers confirmed the impairment of the differentiation process, as evidenced by significantly reduced expression of Keratin 10 and a discontinuous involucrin expression pattern (**Figure 7b**).

Based on these observations, GPRC5A^KD^ RHE were treated with the mimetic polypeptide one day prior exposure to air/liquid interface. Haematoxylin-eosin staining revealed an increase in epidermis thickness along with an augmentation in the number of cell layers (**Figure 7b-c**). The positive effect on epidermal stratification was further confirmed by the restored expression of terminal differentiation markers, including Keratin 10, and to some extent, involucrin (**Figure 7b**). Lastly, transepidermal water loss (TEWL) measurements were conducted to evaluate the inside-out permeability of the RHE barrier function (**Figure 7d**). These measurements indicated a reduction in water loss from the polypeptide-treated GPRC5A^KD^ RHE, with TEWL readings approaching those obtained for control RHE, suggesting a beneficial impact of such treatment on the integrity of the epidermal barrier.

Considering these results, it becomes apparent that the GPRC5A receptor, and more particularly its C-terminal region, is implicated in the regulation of both keratinocyte adhesion and migration, as well as in epidermal differentiation and stratification.

## DISCUSSION

In this study we provide evidences that GPRC5A plays a key role in keratinocyte behavior and epidermis stratification. We indeed demonstrate that GPRC5A expression is upregulated in response to increase in substrate stiffness, with a differential expression of GPRC5A directly linked to the difference between the initial and the final substrate rigidity, thus underlying a potential function of the receptor in keratinocyte mechanosensing.

It is well accepted that during the wound healing process, the mechanical properties of the dermis are disturbed with the formation of the granulation tissue (Hinz, 2010). The sudden rise of ECM stiffness could then participate to the specific location of GPRC5A observed in migrating keratinocytes at the leading edges during reepithelization. It is consistent with previous observations made in mouse wound healing models where GPRC5A displayed similar spatiotemporal expression from day 1 to day 10 post-lesion (Aragona *et al*., 2017). While GPRC5A is repressed once the mouse epidermis is repaired, it remains constitutively expressed in the human differentiated epidermis.

GPRC5A is an orphan receptor and previous studies failed to identify a ligand and its activation remains unclear (Brauner-Osborne *et al*, 2001; Fredriksson *et al*, 2003; Hirano *et al*, 2006). Indeed, the role of GPRC5A in epithelial to mesenchymal transition during cancer development has been intensively studied (Bulanova *et al*, 2017). GPRC5A was shown to play a complex and dual role as oncogene or onco-suppressor depending on the tumor context (Zhou & Rigoutsos, 2014). On one hand, in the lung, GPRC5A high expression at steady-state is required to suppress lung carcinogenesis (Ye *et al*, 2009). Indeed, GPRC5A downregulation in lung cancer is associated to a tumor-suppressive effect by inhibiting Stat3 signaling through Socs3 stabilization (Chen *et al*, 2010). Another group identified GPRC5A as a modulator of epithelial cell proliferation and survival (Zhong *et al*, 2015). Authors described an unexpected direct link between GPRC5A and the translation initiation complex eIF4F that thereby suppresses the translation of membrane-bound proteins (Zhong *et al*., 2015). On the other hand, GPRC5A in several other epithelial tissues is considered as a pro-oncogene via its ability to modulate cell cycle regulation and apoptosis (Zhou & Rigoutsos, 2014). All these observations placed GPRC5A as a tissue-specific regulator responding to cell physical and chemical environments, transducing external signals to modulate cell cycle, adhesion and migration.

Immunofluorescence staining of GPRC5A reveals a dynamic relocation of GPRC5A C-terminal region from cytoplasm to the nucleus during keratinocyte adhesion and differentiation. The distinct feature of having its C-terminal portion cleaved and directed to the nucleus renders GPRC5A a highly unconventional receptor. Presently, only a few transmembrane receptors have been reported to potentially localize within the nucleus, either in their entirety or as cleaved fragments (Ahmad *et al*, 2020; Sprinzak & Blacklow, 2021). Furthermore, in 2006, Cook et al. demonstrated that the angiotensin II type 1 receptor, which is also a GPCR, could undergo cleavage (Cook *et al*, 2006). They observed an accumulation of its C-terminal cytoplasmic portion within the nuclei of endothelial cells. This phenomenon was associated with altered signal transduction and cell proliferation, implying that such cleavage serves as a regulatory mechanism for a biological function. However, to date, no receptor has demonstrated similar activity in the cutaneous context, specifically within keratinocytes. The GPRC5A receptor, which is localized intracellularly and not at the plasma membrane like other GPCRs, appears to function in a manner that is currently undocumented in the literature. While no information currently explains the mechanisms involved in the subsequent translocation of the C-terminal portion of GPRC5A to the nucleus, we identified cathepsin G to be responsible for GPRC5A C-terminal region cleavage. This enzyme is known to be primarily synthesized by immune cells in both secreted and intracellular forms (Schechter *et al*, 1994; Stoeckle *et al*, 2009). It is also found in certain non-myeloid cells, such as endothelial cells, smooth muscle cells, astrocytes, or fibroblasts (Abraham *et al*, 1993; Cavarra *et al*, 2002). As a result, this immune protease has a broader role besides the inflammatory response and is involved in various physiological processes, these include muscle contraction, epithelial renewal or tissue remodeling (Garg *et al*, 2012). Indeed, cathepsin G main function is to cleave intracellular proteins in order to activate them (Dzau *et al*, 1987), but also extracellular matrix proteins to promote immune cell migration (Son *et al*, 2009). Despite no study has reported cathepsin G activity in keratinocytes to date, single-cell analyses conducted on mouse skin have highlighted its expression in the basal keratinocytes of the inter-follicular epidermis (https://kasperlab.org/mouseskin, accessed on 01/10/2023). Furthermore, the results obtained during this work have also demonstrated the expression of this enzyme in primary cultured keratinocytes. Even though verification of cathepsin G expression in whole human epidermis would provide additional support for these findings, we can reasonably assume, based on the co-localization of the enzyme in the Golgi apparatus of keratinocytes (**Figure S9**), that an interaction with the C-terminal region of GPRC5A receptor in the Golgi apparatus lumen is probable. Finally, the inhibition of the enzyme’s activity leading to a significant reduction in the translocation of the receptor’s C-terminal domain into the nucleus also suggests an interaction between the two proteins.

GPRC5A signaling cascade in the skin remains to be fully elucidated, but thanks to the analysis of different kinases phosphorylation, this study brings new evidence about the signaling pathways involved downstream of GPRC5A. Two main pathways were identified to be upregulated by GPRC5A knockdown during the first steps of cell adhesion, namely FAK/JNK and AKT/PRAS40/p53. FAK participates in the formation of focal adhesion between cell and extracellular matrix and mediates an integrin-dependent signal transduction. It was found to be upregulated, phosphorylated, and redistributed to focal adhesion during keratinocytes adhesion (Kim *et al*, 2000), and more particularly during wound healing process (Wang *et al*, 2018), suggesting a functional role of FAK in cell migration. FAK also regulates many signalingpathways, such as PI3K/Akt and MAPK - Erk1/2, JNK - that are closely related to cell proliferation and migration (Mitra & Schlaepfer, 2006; Oktay *et al*, 1999). Our results showed that the inhibition of GPRC5A induced an activation of these pathways, that would result in an exacerbation of keratinocytes adhesion, leading to a slowed migration, which is consistent with our observation showing an increase of cell adhesion in GPRC5A^KD^ cells. More investigations are however needed to better understand the whole mechanism. Indeed, in different epithelial cancer cells, it was shown that GPRC5A inhibition could induce a decrease of adhesion and cell spreading on matrigel, explained by a loss of FAK phosphorylation due to the inhibition of β1-integrin signaling (Bulanova *et al*., 2017). Our observations are consequently in contradiction with these results. In keratinocytes, GPRC5A^KD^ led to an important increase of FAK phosphorylation on the same site (Y397). This difference could be explained by the experimental context since our cells were plated on plastic dishes, while Bulanova’s study used Matrigel^®^ coatings. Another explanation may also rise from the considered cell type – breast cancer epithelial cells or keratinocytes – showing that this receptor can have opposite functions depending on the cell type. AKT is the downstream effector of PI3K with a phosphorylation-dependent activation. It activates the TOR/PRAS40 kinase axis regulating protein synthesis, as well as GSK-3 acting on cell cycle. More precisely, upregulation of the Akt/mTOR signaling pathway was shown to increased oral epithelial cell proliferation and migration, thereby accelerating the wound healing process (Castilho *et al*, 2013). In physiological conditions, mTORC1 regulates MDM2 levels that maintain low p53 protein expression (Havel *et al*, 2015). Because tissue repair involves rapid increases in cell proliferation, it makes sense to expect p53 levels to decrease to allow for the sudden need in increase of cellularity. But hyperactivation of Akt can result in mTORC1-dependent increase in p53 translation in addition to MDM2 sequestration in the nucleolus, inhibiting p53 ubiquitination, leading to a p53 accumulation and to further cellular senescence (Astle *et al*, 2012). In our study, the lack of GPRC5A consequently led to an increased activation of Akt pathway. Specifically, a two-fold increase of p53 phosphorylation on S15 residue, involved in the cellular stress response, was observed. Another clue about cellular stress is the rise of the total protein HSP60 in sh-GPRC5A cells. This mitochondrial protein, in addition to its physiological function under normal, non-stressful conditions also acts as a chaperone when cells are exposed to stress factors. It can besides play a role in regulating apoptosis by interacting directly with key components of the apoptotic pathway and its increase in a context of GPRC5A^KD^ cells, could correspond to a stressful environment. However, whether the lack of GPRC5A could contribute to keratinocyte senescence remains to be demonstrated.

To conclude, we demonstrated that GPRC5A is a novel actor in keratinocyte mechanotransduction during skin wound healing. This receptor appears to be implicated in the regulation of the adhesion and migration pathways, as well as the early stages of the differentiation. These discoveries attest the interest to use this protein as a target to potentialize the reepithelization phase during the skin wound healing. More precisely, by using a polypeptide mimicking the C-terminal region of the receptor, we were able to partially restore a wild-type phenotype, suggesting that this part of the receptor is essential for intracellular signaling. Nevertheless, further functional studies will now be required to identify GPRC5A role in the nucleus.

## MATERIALS AND METHODS

### Ethical considerations

Infant foreskins were collected according to the Declaration of Helsinki Principles. A written informed consent was obtained from infants’ parents according to the French bioethical law of 2004. Abdominal adult skin explants were collected from the Biological Resource Center GCS/CTC (AC-2019-3476, 2^nd^ July 2020) hosted by Hospices Civils de Lyon, in France.

### Cell culture

The human primary epidermal keratinocyte cultures were obtained from child foreskin biopsies or women abdominoplasty after enzymatic treatment as described in (Ya *et al*, 2019). N/TERT-1 cell line was obtained from Rheinwald’s lab (Dickson *et al*, 2000). Keratinocytes were cultivated at 37°C in a 5% CO2 atmosphere in a « keratinocyte serum-free medium (Ker-SFM, GIBCO) » completed with 25 μg/ml bovine pituitary extract, penicillin/streptomycin (100X), 0.2 ng/ml Epidermal Growth Factor and 0.3 mM CaCl2. Presence of mycoplasma in cell cultures was detected regularly using MycoAlert assay (Lonza).

### RNA interference

siRNA targeting human GPRC5A (siGPRC5A_4 against target sequence 5′-CTGGGTGTGTTGGGCATCTTT-3′, cat# SI04438021) and non-targeting control siRNA (cat# Ctrl_AllStars_1, target sequence not disclosed) were purchased from Qiagen. Human primary keratinocytes were transfected at 20 nM final siRNA concentration using Lipofectamin 2000 reagent (Life Technologies) according to the manufacturer’s instructions.

### Generation of polyacrylamide hydrogels

The hydrogels were synthesized according to the principle described previously by (Tse & Engler, 2010). Briefly, 18 mm diameter coverslips were activated with 70 mM NaOH solution and heated at 80°C. This step was repeated with milliQ water until NaOH forms a thin semi-transparent film. The coverslips were treated for 5 min with APES (3-Aminopropyltriethoxysilane, Sigma) and thoroughly rinsed. Then, 0.5% glutaraldehyde (Sigma) was added onto the coverslips for 30 min and dried few minutes at room temperature. In parallel, 24x60 mm glass slides were chlorosilaned with DCDMS (dichlorodimethylsilane, Sigma) for 5 min, gently rinsed, and dried for 30 min at room temperature.

Polyacrylamide gels with different modulus of elasticity values were produced by mixing acrylamide/bis-acrylamide solutions at final percentages (p/v) of 3/0.1, 4/0.225 and 10/0.225 for Soft, Medium and Rigid hydrogels respectively. The polymerization is initiated with APS (ammonium persulfate, Sigma) and TEMED (N,N,N’,N’-tetramethylethane-1,2-diamine, Sigma). The polyacrylamide solution is immediately pipetted onto the chlorosilaned slide and covered by the functionalized coverslip. After the completed polymerization, the top coverslip with the attached polyacrylamide gel is slowly peeled off and rinsed to take out DCDMS on the gel surface.

To facilitate keratinocyte attachment, a heterobifunctional crosslinker, sulfo-SANPAH (sulfosuccinimidyl6(4-azido-2-nitrophenyl-amino)hexanoate, ThermoFisher Scientific), is used to crosslink extracellular matrix molecules onto the surface of the gel. 0.2 mg/mL sulfo-SANPAH solution is added to gel surface, placed 3 inches under an ultraviolet lamp, irradiated for 10 min and rinsed with water. Then, 100 µg/mL of rat tail collagen I solution (ThermoFisher Scientific) is added on the gel and incubated for 2 h at room temperature under a 50 rpm agitation. Hydrogels were rinsed in PBS (phosphate buffered saline), placed in 12-well plates, and immersed in culture media at 37°C one night prior to cell seeding.

### mRNA extraction and qRT-PCR

Total RNA was extracted using RNeasy mini kit (Qiagen, Valencia, CA) according to the manufacturer’s instructions. Reverse transcription was performed using PrimeScript™ RT Reagent Kit (Takara, Tokyo, Japan). qRT-PCR was performed on a Mx3000P real-time PCR system (Stratagene, San Diego, CA, USA) using SYBR® Premix Ex Taq™ II (TaKaRa). Specific primers for RPL13, RPLP0 (ribosomal housekeeping gene) and GPRC5A were used (**Table S1**). The delta delta Ct method was used to calculate the relative fold change.

### RNA sequencing

#### Samples processing

Cells were cultured on a glass substrate and soft hydrogel for 3 days and total RNA was extracted using RNeasy mini kit (Qiagen). RNA quantity and purity were verified using 2200 TapeStation system (Agilent Technologies). Library preparation was performed using mRNA-Seq Library Prep Kit Lexogen following the manufacturer’s instructions. Libraries were validated on TapeStation – HSD1000 ScreenTape®Dosage. Barecoded libraries were pooled together (three per run) on an equimolar basis and run using PI chips on an Ion Torrent™ PGM sequencer using HiQ chemistry. Library preparation and sequencing were achieved by the IGFL sequencing platform (Lyon, France).

#### Bioinformatic analysis

Reads were aligned to the human reference genome hg19 using the Ion Torrent RNASeqAnalysis plugin. Two consecutive alignments were achieved through the STAR and Bowtie2 programs to generate the BAM files. Reads over genes were determined using the R/Bioconductor “Rsubread” software package to create a counts matrix (Liao *et al*, 2013). Differential gene expression analysis was performed using R/Bioconductor “limma” and “edgeR” software packages (McCarthy *et al*, 2012; Ritchie *et al*, 2015; Robinson *et al*, 2010). Heat map were generated from limma-voom normalized values in R (Law *et al*, 2014). Genes with a differential expression FDR ≤ 0.05 were considered significant (**Table S2**).

### Infection with lentiviral particles

Mission ^TM^ lentiviral transduction particles were purchased from Sigma. The pLKO puro lentiviral vector expressed a short hairpin RNA (shRNA) sequence targeting GPRC5A mRNA (shGPRC5A TRCN0000005628) or a control sRNA (shCTR SHC002V). Infection was performed on subconfluent N/TERT-1 cells at a multiplicity of infection of 1, in KGM2 medium supplemented with 1 µg/ml Polybrene (Sigma). Cells that integrated the lentiviral vectors were selected with 1µg/ml Puromycin (Sigma).

### Analysis of kinase’s phosphorylation rate

N/TERT-1 sh-CTRL and sh-GRPC5A cells were plated in 20mm² culture dishes in a complete Ker-SFM medium. After 2 and 24 hours of culture at 37°C, cells were washed with PBS and frozen at −80°C overnight. Levels of protein phosphorylation were then detected using the Human phospho-kinase Array Kit (ARY003B, R&D Systems) according to the manufacturer’s instructions (**Table S3**). Optical density was analysed using ImageJ software for each spot of the array.

### Cell proliferation assays

The cell proliferation rate of transfected keratinocytes with shRNA control or shRNA GPRC5A was measured after 1 to 3 days, using the alamarBlue^TM^ cell viability reagent (Life Technologies). Cells were seeded and the proliferation rate was analyzed by adding 10% of alamarBlue reagent in fresh culture medium and incubated for 4 hours. The fluorescence intensity (excitation 570nm/emission 585nm) was measured using Tecan Infinite M1000 (Life Technologies).

For DNA quantification (Quant-iT™ PicoGreen™ dsDNA Assay Kit), cells were seeded in 96-well plates and DNA concentration was assessed according to the manufacturer’s instructions.

### Cell adhesion assays

#### Percoll assay

Adhesion assay was performed as described in (Goodwin & Pauli, 1995). 96-well plates were coated either with 100 µg/ml of collagen type I (Life Technologies) or 1% bovine serum albumin (BSA) in phosphate buffered saline (PBS) (negative control). Cells were trypsinized with 10 mM EDTA for 10 min at 37°C and washed twice with serum-free medium. Then 2.5×10^4^ cells/well were re-suspended in a serum-free medium and plated in 5 replicas for each condition. Cells were allowed to attach for 5, 10, 30, 60, 120 and 240 min at 37°C, then unattached cells were removed with Percoll solution containing 0.57 mM NACl. The remaining cells were fixed with Peroll solution containing 5% of glutaraldehyde for 15 min at room temperature. The cells were stained with 0.1% Crystal Violet for 30 min at room temperature, and then the plates were washed with water tap. Plates were air dried, and the crystal violet was then extracted using 2% SDS in distilled water for 30 min on a shaker. Absorbance at 570 nm was measured on Tecan Infinite M1000 (Life Technologies). Background absorbance (from blank wells) was subtracted from all test wells. The assay was always performed in 5 technical and 3 biological replicas.

#### Impedance measurement

Impedance of cells was measured using the xCELLigence system, as described in [Hamidi et al. – 2017]. Briefly, E-plates were coated with 100 µg/ml of Collagen type I and then saturated with 0,1% BSA (w/v). Cells were pre-treated or not with 10 µg/ml of mimetic peptide, during 24 hours before being harvested and replated in the E-plate (20 000 cells/well in 100 µl) in 4 technical replicates. Impedance is then measured for at least 5 hours after plugging the E-plate in the xCELLigence system placed inside the incubator at 37°C.

### Cell migration assays

#### Ibidi insert

In 6-well plate, cells (2×10^4^ cells/well) were seeded in 2-well silicone insert (Ibidi) for 30 hours. After confluence, the culture insert was slowly removed. After washing with PBS, the cells were supplied with a 2 ml culture medium. The cells were then allowed to migrate into the gap (500 µm) after 12, 14, 16, 18, 20, 22 and 24 hours. Quantification of cell migration was performed by measuring uncovered areas using Image J software. Each experiment was repeated in 3 technical and 3 biological replicas.

#### Scratch assay

Cells were seeded in 24-well plate (2,5×10^5^ cells/well), beforehand coated with 100 µg/ml of Collagen type I. After 24 hours of adhesion, cells were then either treated or not with 10 µg/ml of mimetic peptide for another 24 hours. Then a scratch is performed using some 20 µl tip, the cells are rinced with PBS, fresh medium is added, and the closure of the wound is recorded using video microscopy (Zeiss Axio Observer 7) for 15 hours.

### Western blot

Proteins were extracted in RIPA buffer (50 mM Tris-HCl, pH 8, 150 mM NaCl, 1% Nonidet P-40, 0.1% sodium deoxycholate, 0.1% SDS, 1 mM orthovanadate) and protease inhibitor cocktail (Thermo Fischer scientific). Lysates were centrifuged for 10 min at 14000 g at 4°C to eliminate cell debris. Protein concentration was finally measured using BCA assay (ThermoFischer). They were separated by SDS–PAGE followed by transfer to polyvinylidene fluoride membrane (EMD-Millipore, Billerica, MA). The membrane was blocked with 5 % non-fat milk in Tris buffered saline (TBS) buffer containing 0.1% Tween-20, and incubated with rabbit anti-GPRC5A antibody (HPA007928, Sigma, 1:250), mouse anti-actin (C4, MAB1501, Sigma-Aldrich, 1:5000), rabbit anti-akt (9272, CellSignaling, 1:1000), rabbit anti-Phospho-akt (SER473) (9277S, CellSignaling, 1:1000) and rabbit anti-cathepsin G (703590, Invitrogen, 1:1000), overnight at 4°C. The membrane was incubated with secondary antibodies for 1 h at room temperature: goat anti-rabbit IgG (H+L)-HRP conjugate (170-6510, Biorad, 1:10000) and goat anti-mouse IgG (H+L)-HRP conjugate (170-6516, Biorad, 1:10000). Antibody binding was detected by the enhanced chemiluminescence system (Thermo Fischer Scientific), using a Fusion Fx system (Vilber Lourmat).

### Preparation of reconstructed epidermis

The reconstructed epiderma were produced according to the principle described previously by (Le Provost *et al*, 2010). Briefly, fibroblasts (2.5×10^4^) were seeded underneath the polycarbonate membrane of the cell culture insert (Nunc, ThermoFischer Scientific). After 4 hours, the inserts were placed in 24-well plates, and fibroblasts were cultured for 2 days in DMEM/F12 (1/1) supplemented with 10 % fetal bovine serum (Fetal Clone II; Hyclone, Thermo Scientific, France) and 1% Penicillin/Streptomycin (Sigma). Then, N/TERT-1 infected by pLKO shRNA control or pLKO shRNA GPRC5A (2.5×10^5^) were seeded on the top of the membrane and cultured for 3 days in DMEM/F12 (1/1) supplemented with 5 % fetal bovine serum (Fetal Clone II; Hyclone, Thermo Scientific, France), 0.2 ng/ml EGF (Gibco), 0.4 µg/ml Hydrocortisone (Sigma), 5 µg/ml Insulin (Sigma), 8 ng/ ml Cholera Toxin (Sigma), 2×10^-11^ M Tri-iodothyronine (Sigma), 24 µg/ml Adenine (Sigma) and 1% Penicillin/Streptomycin. 24 hours the air/liquid interface, knock-down keratinocytes were treated or not with 10 µg/ml of mimetic peptide. To induce stratification and differentiation, keratinocytes were then placed at the air/liquid interface and cultured for 2 days with the same medium, except that EGF and adenine were omitted, and the final concentration of calcium chloride (Sigma) was adjusted to 2 mM and 50 µg/ml vitamin C. Cells were then cultured for another 14 days in the same medium, except that the fetal bovine serum was reduced to 1%. During the emersion phase, the culture medium was changed every day.

### Functional evaluation of barrier function

Trans-epidermal water loss (TEWL) was measured using a Tewameter (TM 300) on 16 days RHE. The probe measures the density gradient of the water evaporation from the skin (in g/h/m²) indirectly by the two pairs of sensors (temperature and relative humidity) inside the hollow cylinder.

### Human skin ex vivo injury model

A heated metal piece (2mm width, 150°C) was applied at the center of the skin sample for 3 seconds. Burned skins were then cultivated for 12 days on culture inserts to maintain the epidermis at the air/liquid interface. Skins were cultured in DMEM/F12 medium (Gibco) supplemented with Zellshield antibiotic (13-0150, Clinisciences).

### Histology and immunofluorescence microscopy

RHE, human skin and cells on coverslip were fixed in 4% paraformaldehyde (PFA) solution, for 24 hours at 4°C. Samples were then embedded in paraffin and cut into 5 µm sections. After dewaxing and rehydration, tissue sections were stained with hematoxylin and eosin solutions for routine histology or permeabilized by 0.1% triton X100 for 10 min and then boiled in 10 nM citrate buffer pH6 for antigen retrieval. For immunocytofluorescent labelling, cells on coverslip were fixed for 10 minutes in 4% PFA solution. Then cells were permeabilized by 0.1% triton X100 for 10 minutes. Both skin sections and cells were then blocked by 5% of bovine albumin serum and incubated with the following primary antibodies overnight at 4°C. Cells were finally incubated with primary antibodies overnight: rabbit anti-GPRC5A (HPA007928, Sigma Aldrich, 1:250); rabbit anti-cytokeratin 10 [EP1607IHCY] (ab76318, Abcam, 1:500); mouse anti-cytokeratin 14 (MA5-11599, Invitrogen, 1:500), rabbit anti-involucrin (ab53112, Abcam, 1:500) and rabbit anti-cathepsin G (703590, Invitrogen, 1/100). Secondary antibodies were: alexa-488 and alexa-546 -conjugated goat anti-mouse IgG or anti-rabbit IgG (Molecular Probes, Life Technologies). Nuclear counterstaining using DAPI (4’,6-diamino-2-phenylindole) (Sigma) was carried out. Sections and coverslip were then mounted in ProLong™ Gold Antifade Mountant (Invitrogen). Image acquisition was performed using a fluorescent TiE Nikon microscope or a Leica SP5X confocal microscope, in the CIQLE platform “Centre d’imagerie Quantitative Lyon-Est de l’Université Lyon I”.

### GPRC5A translocation kinetic

Primary keratinocytes were cultivated on soft polyacrylamide hydrogels (4kPa) for 3 days, then cells were transferred on stiff support (glass slide). During the transfer, keratinocytes were either treated or not with a Cathepsin G inhibitor (219372 – Sigma) at a concentration of 10 µM for 30 minutes. Depending on the experiment, GPRC5A location was observed after 30 minutes, 1, 4 and 24 hours, but also after cells were confluent and 5 days post-confluency. Two different antibodies were used to label GPRC5A: rabbit anti-GPRC5A (HPA007928, Sigma Aldrich, 1:250) targeting the C-terminal region of the receptor, and mouse anti-RAI3 (PA5-28738, Invitrogen, 1:250) targeting its N-terminal region. Intracellular compartment labelling was also performed using mouse anti-nucleolin (Ab154028, Abcam, 1/1000), concanavalin (C11252, Invitrogen, 1:100) and mouse anti-Golgi 58k (MA1-22144, Invitrogen, 1/250).

### Mimetic C-ter peptide production and purification

GPRC5A C-terminal domain sequence was amplified from cultured keratinocytes and cloned into pET30a plasmid digested by EcoRV. Then a poly(R/K) tag was added by amplifying the C-ter sequence with some primer containing the tag, and the TEV-6His sequence was extracted from a pT7 plasmid to form a final plasmid pET30a_GPRC5A-C-terminal_poly(R/K)_TEV-cleavage-site_2x-6Histidines (**Table S4**).

BL21 bacteria were transformed with an overexpressing recombinant plasmid containing the C-terminal region of GPRC5A. The polypeptide was produced by adding 1 mM of IPTG in HYPER BROTH (AthenaES AE-0107) after bacteria arrived in the exponential growth phase. After 6 hours of culture, bacteria were centrifugated and protein were extracted: the soluble proteins with 500 mM NaCl, 10 mM MgCl2 and 0.5% Triton X-100 solution, and then the insoluble part with 500 mM NaCl and 8 M urea solution.

Bacteria BL21 (Life Tech 44-048) were transformed with the plasmid pET30a_GPRC5A-C-terminal_poly(R/K)_TEV-cleavage-site_2x-6Histidines and subsequently cultured on LB-agar plates supplemented with kanamycin. A preculture was then initiated, where a single bacterial colony was inoculated into 100 mL of LB BROTH LENNOX medium (Formedium LBX0102) and incubated overnight at 37°C with agitation at 150 rpm. From this preculture, bacteria were inoculated to an optical density (D.O) of 0.15 in HYPER BROTH medium (AthenaES AE-0107) and cultivated at 37°C with agitation at 150 rpm. The growth curve was monitored by measuring the D.O hourly to track growth (ln(DO) over time). During the exponential growth phase of the bacteria (D.O=0.8-1), 1 mM IPTG (Isopropyl β-D-1-thiogalactopyranoside) was added to the culture medium to induce the production of the polypeptide. The culture was then continued until reaching the stationary phase. Subsequently, bacterial pellets were harvested by centrifugation (4000g, 10 minutes at 4°C), separated from the supernatant, and frozen at −20°C.

For protein extraction, bacteria were lysed in a lysis buffer (PBS1X, 500 mM NaCl, 10 mM MgCl2, 0.5% Triton X100 – volume equal to 1/10th of the bacterial culture volume). The lysis buffer was supplemented with protease inhibitors (cOmplete™ Protease Inhibitor Cocktail, Roche), DNAse (100 µg/mL), and lysozyme (10 mg/mL). The lysate was incubated for 30 minutes on a rotator at 4°C and then subjected to sonication (2 rounds of 5 minutes, pulse 3 seconds, amplitude 30). Cellular debris and the insoluble fraction were separated from the soluble fraction by centrifugation (15000g, 15 minutes at 4°C). The soluble fraction was stored at 4°C, while the insoluble fraction was resuspended in a urea buffer (PBS1X, 8M urea, 500 mM NaCl, volume equal to 1/4th of the lysis buffer volume). This resuspended fraction was supplemented with DNAse (100 µg/mL) and incubated for 1 hour on a rotator at 4°C. Proteins released into the buffer were collected in the supernatant by centrifugation (15000g, 15 minutes at 4°C) and combined with the previously saved soluble fraction.

The entire polypeptide was subsequently purified by affinity chromatography using Ni-NTA beads (745400 Propino, Machery-Nagel, volume equal to 1/1000th of the bacterial culture volume). The beads, pre-washed with PBS1X, were added to the protein suspension and incubated for 1 hour on a rotator at 4°C. The beads were then loaded onto a column (with a cotton filter that did not exceed the bead size), and the non-retained fraction was removed by aspiration using a peristaltic pump (1 mL/minute). The beads were further washed with 1M NaCl bead volume, followed by five volumes of a washing buffer (PBS1X, 500 mM NaCl, 10 mM MgCl2, 15 mM imidazole).

Subsequently, the polypeptide was eluted from the Ni-NTA beads by adding an elution buffer (PBS1X, EDTA 50 mM, volume approximately 10 times that of the bead volume) and quantified using the Pierce™ BCA Protein Assay Kit. Without delay, the polypeptide in solution was supplemented with 1 mM DTT (dithiothreitol) and digested by adding TEV protease (Produced by the Protein Science Facility – SFR Bioscience – 0.5 mg TEV for 10 mg of protein). The mixture was placed in a dialysis bag rinsed with PBS1X (MWCO: 3500 Da) and dialyzed against a solution of PBS1X and 500 mM NaCl overnight at 4°C. The cleaved peptide was finally purified by adding Ni-NTA beads as previously described (1 mL of beads per 50 mg of peptide) for 1 hour on a rotator at 4°C. After placing the peptide-bead suspension on an identical column as before, the cleaved peptide was recovered in the non-retained fraction, quantified, aliquoted, and frozen at −80°C.

### N-TAILS sample set-up and mass spectrometry parameters

N-TAILS experiment was used to identify the free amines on the N-terminal part of the GPRC5A cleaved part by mass spectrometry. N/TERT-1 keratinocytes were transfected with an overexpression plasmid containing the C-terminal region of GPRC5A linked to a GFP tag (see the sequence in **Table S5**). Subconfluently cells were transfected using 2 µg/ml of plasmid and Lipofectamin 2000 reagent (Life Technologies) according to the manufacturer’s instructions. After 24 hours, total proteins were extracted and labelled with some isobar TMTsixplex™ kit following the manufacturer’s instructions (Weng *et al*, 2019).

Briefly, the six samples (160 µg of total protein per condition) were dissolved in 180 mM HEPES pH 8 and reduced with 10 mM Tris (2-carboxyethyl) phosphine, alkylated with 25 mM iodoacetamide. Samples were cleaned and concentrated by using SP3 beads (Hughes *et al*, 2019) before labeled with one TMT label (126, 127N; TMT 10-plex kit 90110 from Thermo Scientific), dissolved in DMSO in a 1:5 (total protein/TMT label) mass ratio for 60 min. Labeling reactions were stopped by incubation with 5% hydroxylamine (Sigma) for 30 min. Before digestion, all sample were combined in Hepes 180mM and trypsin was added (trypsin/total protein (1:100); Trypsin V511A, Promega) overnight at 37°C. N-terminal peptide enrichment was performed on the digested sample by removing the internal tryptic peptides with undecanal (Sigma, U2202). Enriched N-terminal peptides were then desalted with a C18 spin column (Thermo Fisher Scientific) and the eluate fraction was freeze-dried, resuspended in 0.1% formic acid and analyzed by LC-MS/MS on a Q-Exactive HF mass spectrometer (three replicates), as described for ATOMS experiments. Data files were analyzed with Proteome Discover 2.4 using the SEQUEST HT algorithm against the human protein database (SwissProt release 2019-12, 43835 entries). Precursor mass tolerance and fragment mass tolerance were set at 10 ppm and 0.02 Da, respectively, and up to 2 missed cleavages were allowed. Oxidation (M, P), pyroglutamate N-term (Q, E), acetylation (Protein N-terminus), and TMT6Plex (N-term, K) were set as variable modifications, and carbamidomethylation (C) as fixed modification.

### Cathepsin G digestion

*In vitro* digestions were performed using purified Cathepsin G to prove the ability of the protease to digest the C-terminal region of the GPRC5A receptor. 3µg of purified mimetic C-ter polypeptide produced in bacteria were submitted to 0.001 unity of Cathepsin G (C4428, Sigma Aldrich) for 1 or 2 hours, at 37°C. Then digested peptides were analysed on polyacrylamide gel, stained with Coomassie thereafter.

### Statistical analysis

All data were presented as mean ± SD for three independent experiments. Statistical significance (p< 0.05) was determined by performing either an unpaired T-test when comparing two groups or one-way or two-way analysis of variance (ANOVA) followed by a Bonferroni post-test when comparing multiple groups. All analyses were performed with GraphPad Prism5 software.

## DATA AVAILABILITY STATEMENT

The data that support the findings of this study are available on request from the corresponding author (RD).

RNA-Seq data: Gene Expression Omnibus GSE248355 (https://www.ncbi.nlm.nih.gov/geo/query/acc.cgi?acc=GSE248355).

The mass spectrometry proteomics data have been deposited to the Center for Computational Mass Spectrometry repository (University of California, San Diego) via the MassIVE tool with the dataset identifier MassIVE MSV000093119.

## ACKNOWLEDGEMENTS

This work was supported by Isispharma France. Choua Ya is a recipient of a PhD grant from the French National Association of Research and Technology (ANRT). Sarah Chanteloube is supported by a PhD grant from the French ministry of Higher Education and Research.

We acknowledge the contribution of Catherine Moali but also the SFR Biosciences (UMS3444/CNRS, US8/Inserm, ENS de Lyon, UCBL) facilities: the staff of PSF (Protein Science Facility), especially Frédéric Delolme and Adeline Page, for their help in analyzing the mass spectrometry experiment; the IGFL’s sequencing platform for RNA-seq library preparation and sequencing.

We acknowledge the contribution of SFR Santé Lyon-Est (UAR3453 CNRS, US7 Inserm, UCBL) facilitiy : CIQLE (a LyMIC member), especially Denis Ressnikoff, Bruno Chapuis, Annabelle Bouchardon and Smatti Batoule for their help in confocal microscopy acquisition and histology sample preparation.

## AUTHOR CONTRIBUTIONS

<u>Conceptualization:</u> RD, BF, FN

<u>Formal Analysis:</u> SC, CY, RD

<u>Funding acquisition:</u> RD, BF

<u>Investigation</u>: SC, CY, RD, AB, CD, GLP

<u>Methodology</u>: SC, CY, RD, GLP

<u>Project Administration:</u> RD, BF

<u>Supervision:</u> RD

<u>Visualization:</u> SC, CY, RD

<u>Writing – Original Draft Preparation:</u> SC, CY, RD

<u>Writing – Review and Editing:</u> SC, CY, GLP, CD, SVLG, BF, RD.

## Abbreviations

GPRC5A: G-Protein Coupled Receptor, Class C, Group 5, Member A
N-TAILS: N-terminal amine isotopic labeling
ECM: Extracellular matrix
AFM: Atomic force microscopy
GPCRs: G protein-coupled receptors
RAI3: Retinoic acid-induced protein 3
MS: Mass spectrometry
LC: Liquid phase
KD: Knock-down
RHE: Reconstructed human epidermis
TEWL: Transepidermal water loss

