## Supplementary Figures for "GPRC5A regulates keratinocyte adhesion and migration through nuclear translocation of its C-terminus region"

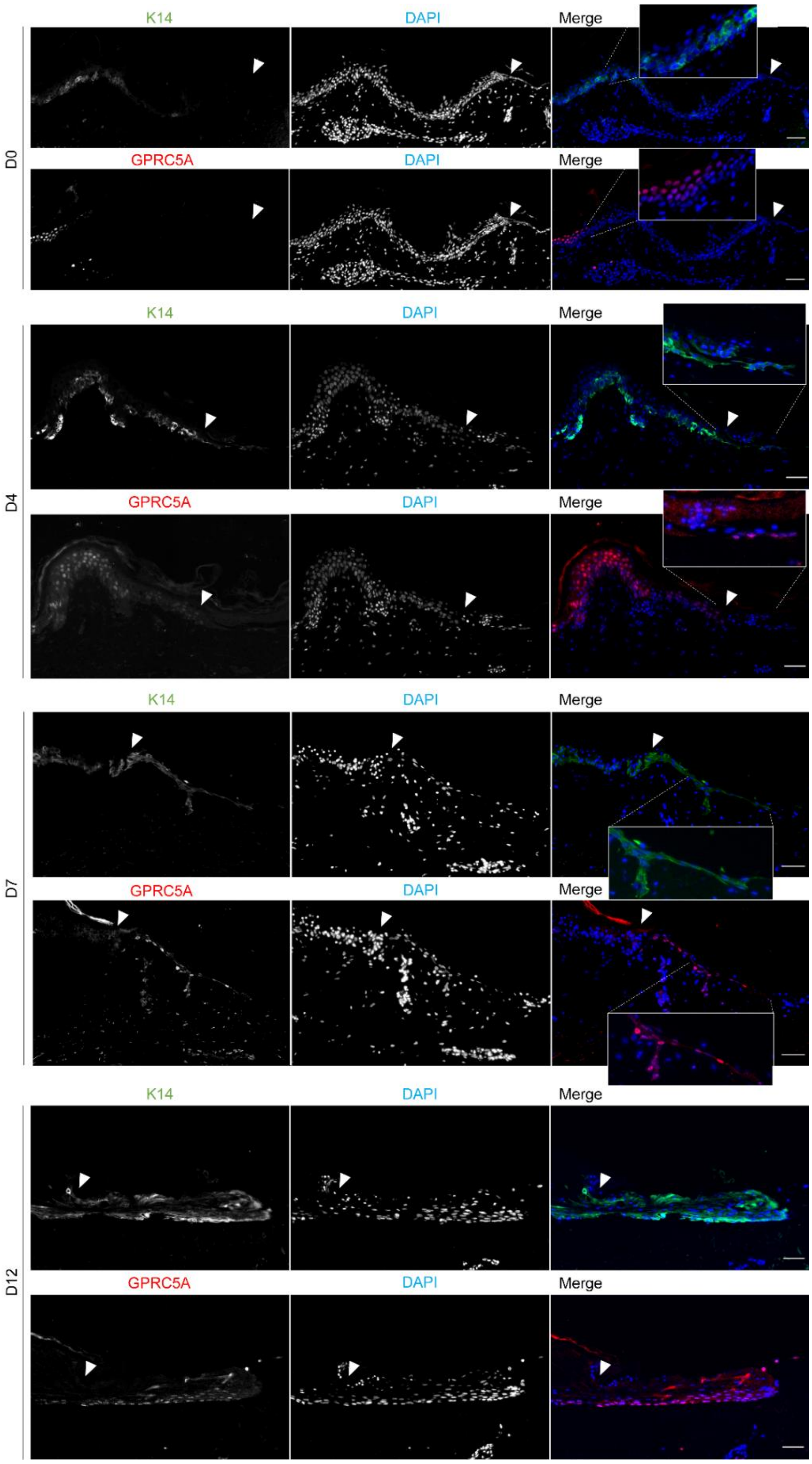


**<u>Figure S1:</u>** GPRC5A expression in ex vivo wound healing of human skin. GPRC5A (red) and Keratin 14 (green) immunostaining after 0-, 4-, 7- and 12-days post-burn (150°C, 3 seconds) in ex vivo abdominal skin. Nucleus were counterstained with DAPI (blue). The white arrow stands for the edge of the wound. Scale bars 25 µm.

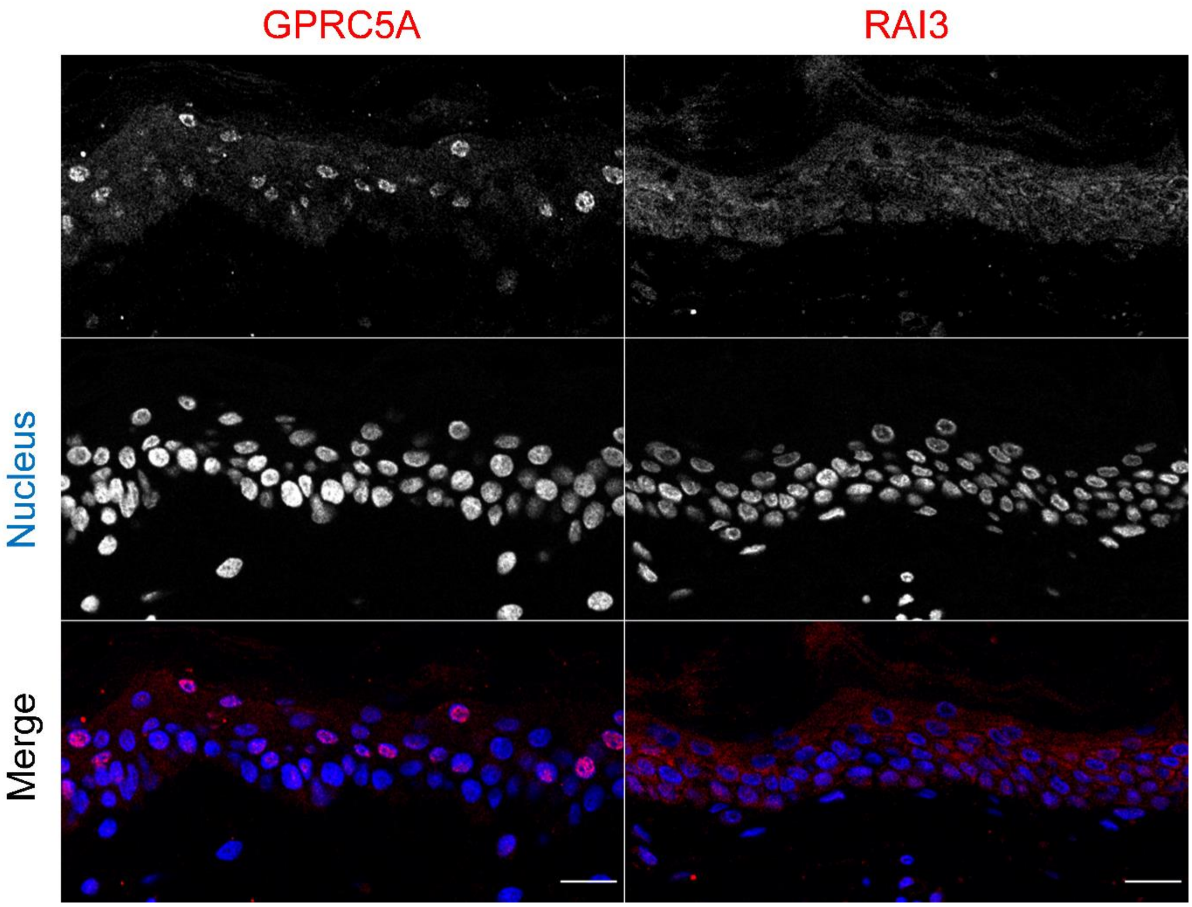


**<u>Figure S2:</u>** GPRC5A immunostaining (red) with anti-GPRC5A (C-terminal part of GPRC5A) or anti-RAI3 (N-terminal part of GPRC5A) antibody, and nucleus (blue) in human abdominal skin. Scale bars: 20µm.

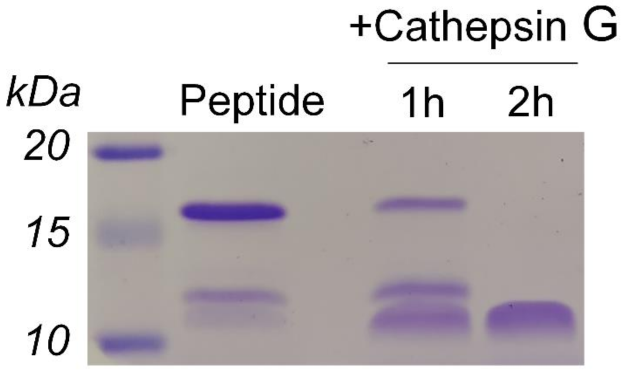


**<u>Figure S3:</u>** Coomassie gel staining of the C-terminal region of GPRC5A mimetic purified polypeptide, either digested or not with Cathepsin G for 1 or 2 hours.

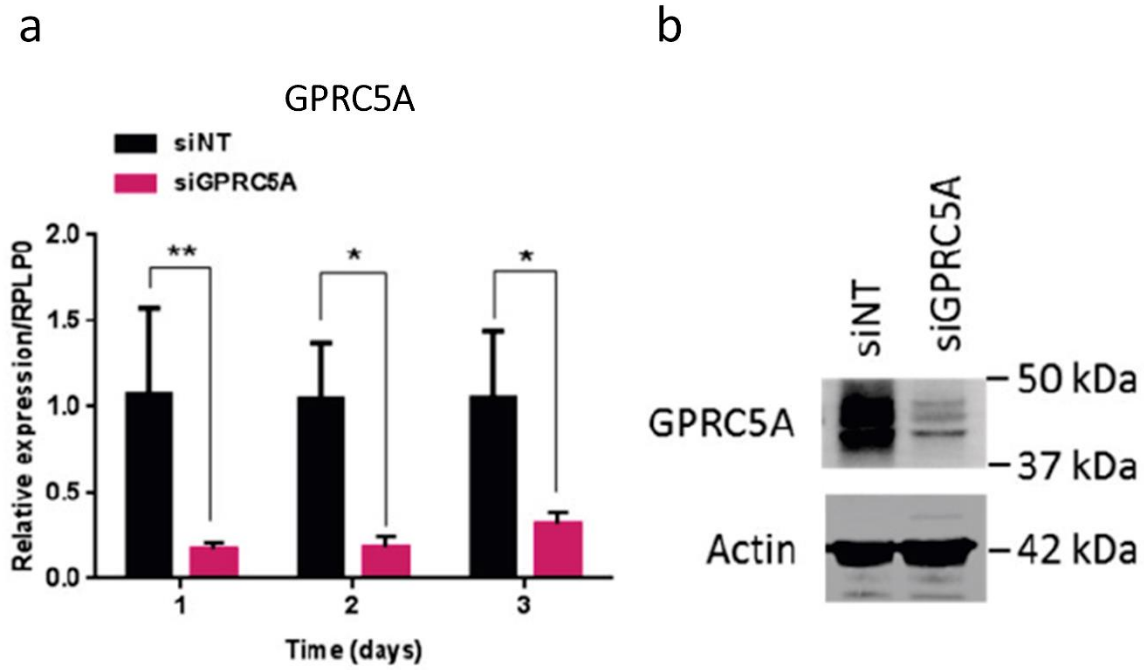


**<u>Figure S4:</u>** Validation of GPRC5A inhibition by siRNA in human primary keratinocytes by qRT-PCR (a) and western blot (b) in human keratinocytes. One-way ANOVA test; * P< 0.05 and ** P<0.01.

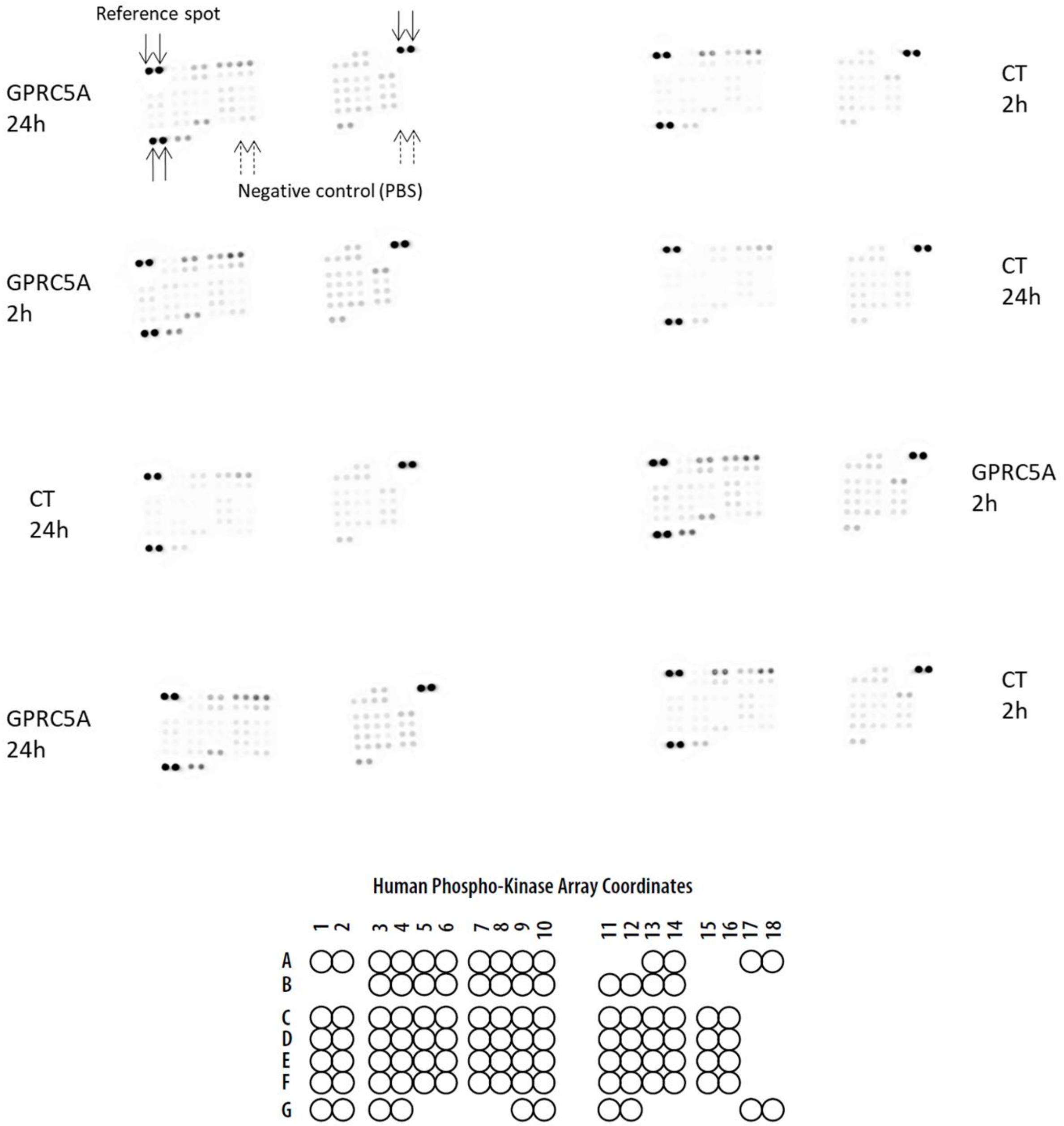


**<u>Figure S5:</u>** Human phospho-kinase array coordinates (by R&D Systems). See legend of the spot position in **Table S3**.

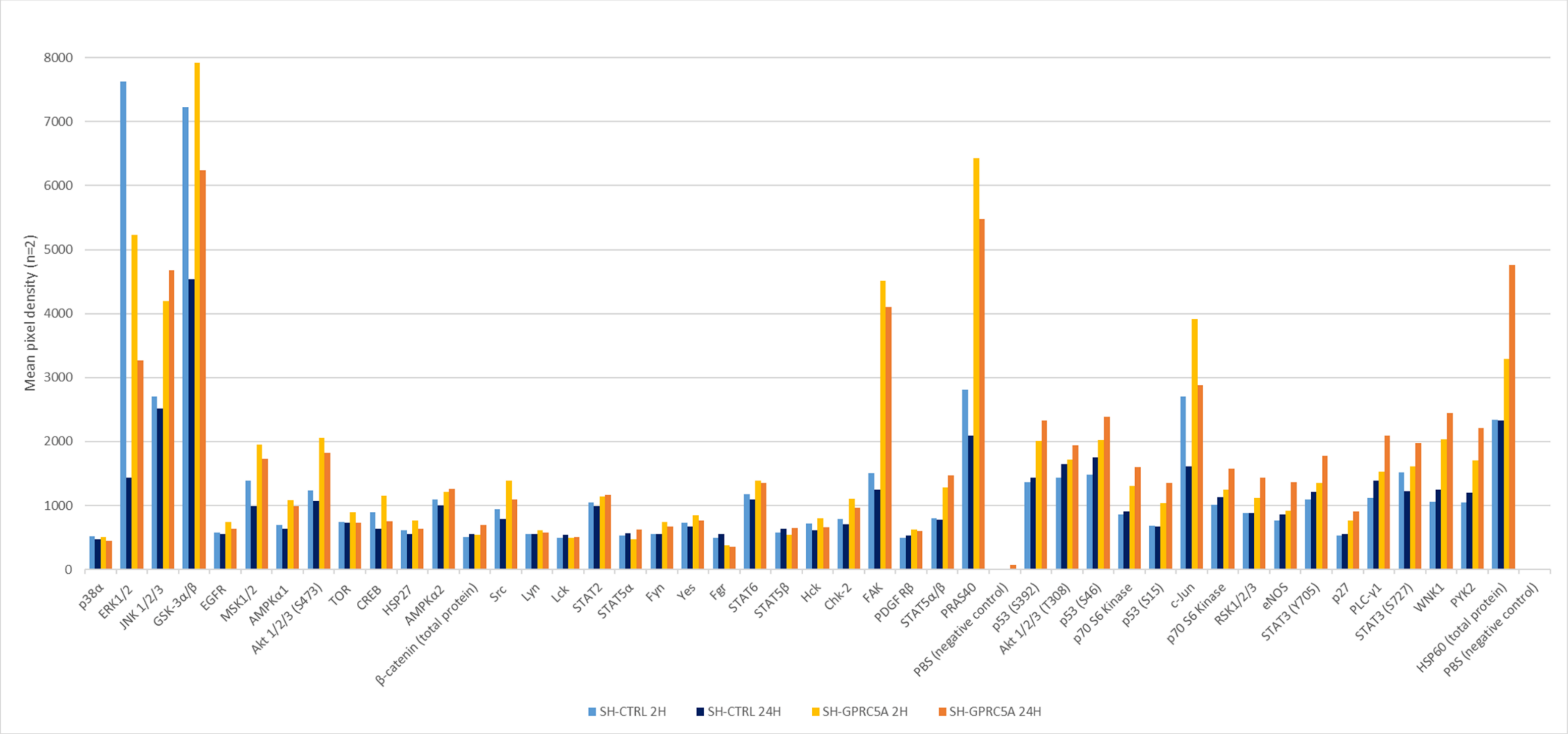


**Figure S6:** Normalized data analysis of kinases phosphorylation in NTERT/1 sh-GPRC5A vs sh-CTRL cells at 2 and 24hours post-adhesion (n=2).

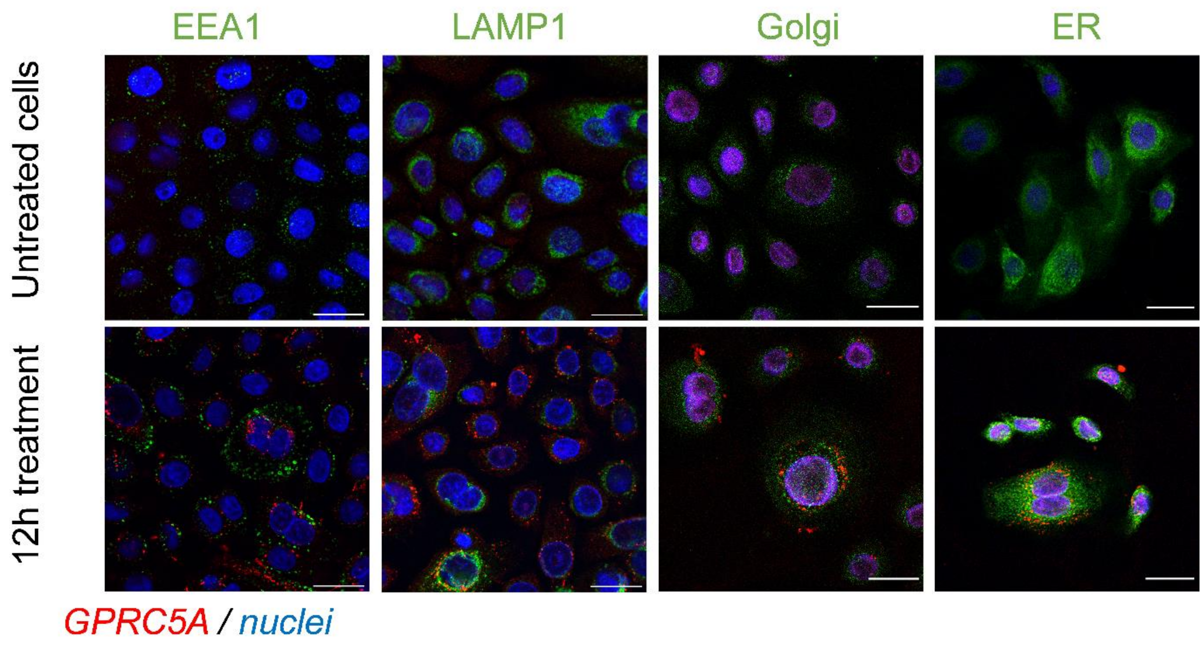


**<u>Figure S7</u>*:*** Fluorescent immunostaining of NHEK1 cells treated with 10 µg/ml of peptide for 12h and observed by confocal microscopy. GPRC5A (red), cellular compartment (green): early endosomes (EEA1), lysosomes (LAMP1), Golgi apparatus, endoplasmic reticulum (concA), and the nuclei (blue). Scale bars : 25 µm.

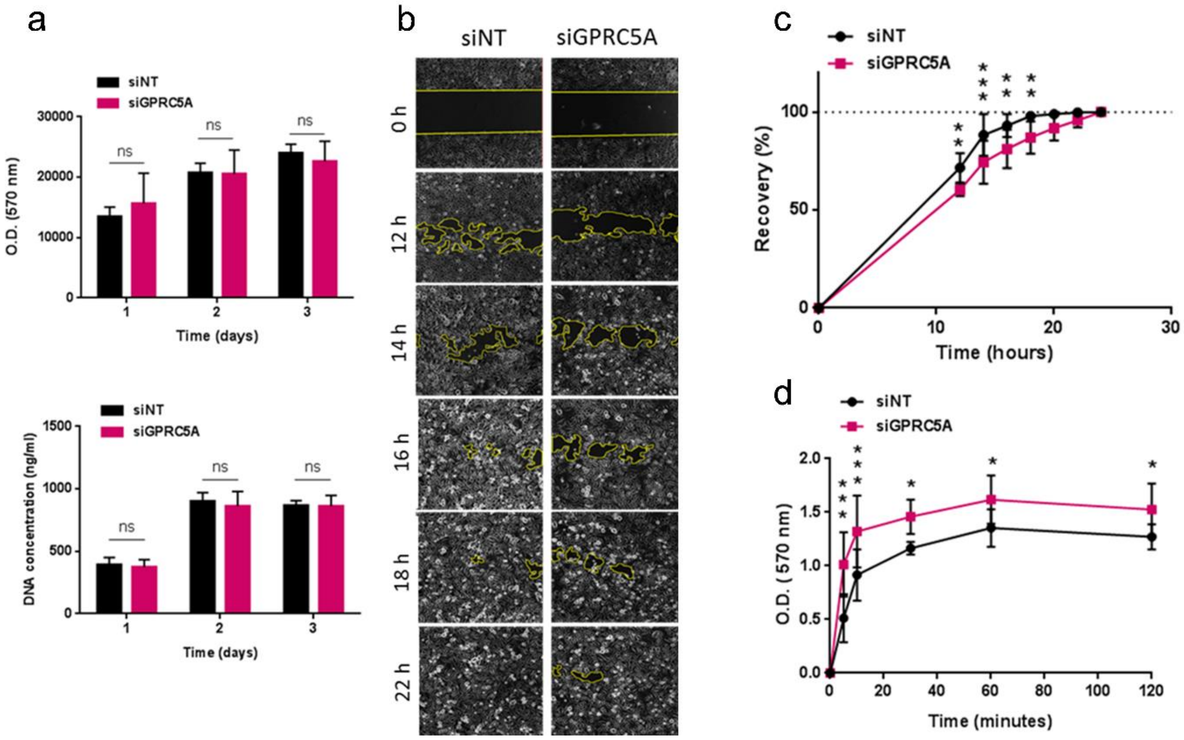


**Figure S8:** Knockdown of GPRC5A in NHEK cells. (a) Proliferation assays. One-way ANOVA test. (b-c) quantification of migration assay. Yellow line represents uncovered area. (d) Quantification of adhesion assay. Two-way ANOVA test; * P< 0.05, ** P<0.01 and *** P<0.001. All results are presented as means ± SD and were performed in at least three independent biological replicates.

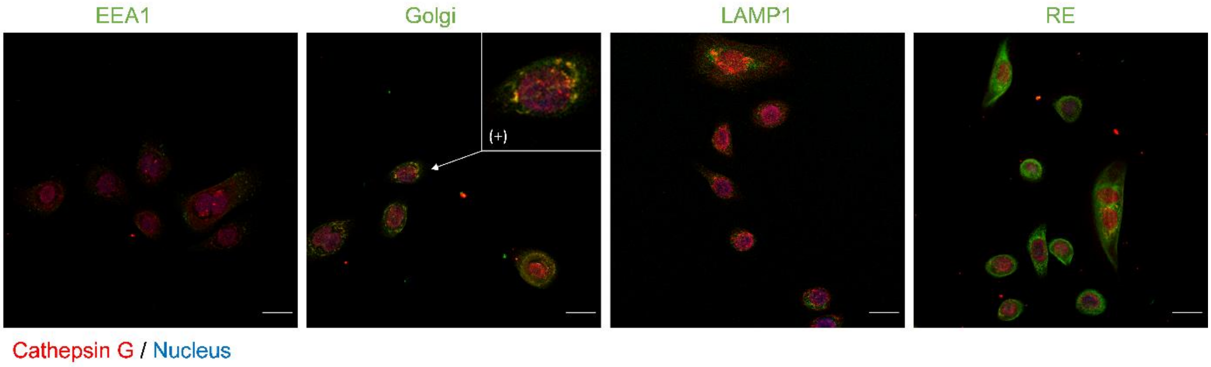


**Figure S9:** Fluorescent immunostaining of Cathepsin G (red), nuclei (blue) and cellular compartments (green) : early endosome (EEA1), Golgi apparatus (Golgi), Lysosome (LAMP1) and the reticulum endoplasmic (RE). Scale bare 20 µm. The zoom in (+) zone shows the colocalization between the Cathepsin G in the Golgi apparatus.
