## Supplementary Tables for "GPRC5A regulates keratinocyte adhesion and migration through nuclear translocation of its C-terminus region"

**Table S1**: Real time PCR primer sequence.

| Gene | Primer forward | Primer reverse |
|---|---|---|
| RPL13A | GATGGTGGTTCCTGCTGCCC | GGCTTTCTCTTTCCTCCTCTCCTCC |
| RPLP0 | CCATTCTATCATCAACGGGTACAA | TCAGCAAGTGGGAAGGTGTAATC |
| GPRC5A | TCTCAAGAGGAAATCACTCAAGGT | GTGGGATGGAGAATTCCTTTT |

**Table S2:** List of gene significantly up- and down-regulated in Soft_Glass conditions (FDR q-value < 0.05).

Up-regulated genes:

| ENTREZID | SYMBOL | GENENAME |
|---|---|---|
| 148304 | C1orf74 | chromosome 1 open reading frame 74 |
| 3664 | IRF6 | interferon regulatory factor 6 |
| 84665 | MYPN | myopalladin |
| 6712 | SPTBN2 | spectrin beta, non-erythrocytic 2 |
| 9052 | GPRC5A | G protein-coupled receptor class C group 5 member A |
| 693199 | MIR614 | microRNA 614 |
| 3854 | KRT6B | keratin 6B |
| 3853 | KRT6A | keratin 6A |
| 479 | ATP12A | ATPase H+/K+ transporting non-gastric alpha2 subunit |
| 8796 | SCEL | sciellin |
| 60485 | SAV1 | salvador family WW domain containing protein 1 |
| 56204 | FAM214A | family with sequence similarity 214 member A |
| 400451 | FAM174B | family with sequence similarity 174 member B |
| 9235 | IL32 | interleukin 32 |
| 283932 | FBXL19-AS1 | FBXL19 antisense RNA 1 (head to head) |
| 374286 | CDRT1 | CMT1A duplicated region transcript 1 |
| 3860 | KRT13 | keratin 13 |
| 201294 | UNC13D | unc-13 homolog D |
| 7077 | TIMP2 | TIMP metallopeptidase inhibitor 2 |
| 1830 | DSG3 | desmoglein 3 |
| 374918 | IGFL1 | IGF like family member 1 |
| 5655 | KLK10 | kallikrein related peptidase 10 |
| 43847 | KLK14 | kallikrein related peptidase 14 |
| 162963 | ZNF610 | zinc finger protein 610 |
| 11227 | GALNT5 | polypeptide N-acetylgalactosaminyltransferase 5 |
| 150094 | SIK1 | salt inducible kinase 1 |
| 9582 | APOBEC3B | apolipoprotein B mRNA editing enzyme catalytic subunit 3B |
| 79443 | FYCO1 | FYVE and coiled-coil domain containing 1 |
| 56999 | ADAMTS9 | ADAM metallopeptidase with thrombospondin type 1 motif 9 |
| 389136 | VGLL3 | vestigial like family member 3 |
| 64393 | ZMAT3 | zinc finger matrin-type 3 |
| 200958 | MUC20 | mucin 20, cell surface associated |
| 10769 | PLK2 | polo like kinase 2 |
| 6584 | SLC22A5 | solute carrier family 22 member 5 |
| 389337 | ARHGEF37 | Rho guanine nucleotide exchange factor 37 |
| 1832 | DSP | desmoplakin |
| 154091 | SLC2A12 | solute carrier family 2 member 12 |
| 6446 | SGK1 | serum/glucocorticoid regulated kinase 1 |
| 79660 | PPP1R3B | protein phosphatase 1 regulatory subunit 3B |
| 63898 | SH2D4A | SH2 domain containing 4A |
| 203111 | ERICH5 | glutamate rich 5 |
| 79987 | SVEP1 | sushi, von Willebrand factor type A, EGF and pentraxin domain containing [illegible] |

| | | |
|---|---|---|
| 9128 | PRPF4 | pre-mRNA processing factor 4 |

Down-regulated genes:

| ENTREZID | SYMBOL | GENENAME |
|---|---|---|
| 54206 | ERRFI1 | ERBB receptor feedback inhibitor 1 |
| 79180 | EFHD2 | EF-hand domain family member D2 |
| 3925 | STMN1 | stathmin 1 |
| 114784 | CSMD2 | CUB and Sushi multiple domains 2 |
| 260293 | CYP4X1 | cytochrome P450 family 4 subfamily X member 1 |
| 51668 | HSPB11 | heat shock protein family B (small) member 11 |
| 9635 | CLCA2 | chloride channel accessory 2 |
| 84722 | PSRC1 | proline and serine rich coiled-coil 1 |
| 26191 | PTPN22 | protein tyrosine phosphatase, non-receptor type 22 |
| 9507 | ADAMTS4 | ADAM metallopeptidase with thrombospondin type 1 motif 4 |
| 57559 | STAMBPL1 | STAM binding protein like 1 |
| 8519 | IFITM1 | interferon induced transmembrane protein 1 |
| 7070 | THY1 | Thy-1 cell surface antigen |
| 9638 | FEZ1 | fasciculation and elongation protein zeta 1 |
| 64581 | CLEC7A | C-type lectin domain family 7 member A |
| 7786 | MAP3K12 | mitogen-activated protein kinase kinase kinase 12 |
| 3221 | HOXC4 | homeobox C4 |
| 11247 | NXPH4 | neurexophilin 4 |
| 10631 | POSTN | periostin |
| 100506411 | LOC100506411 | uncharacterized LOC100506411 |
| 10509 | SEMA4B | semaphorin 4B |
| 9143 | SYNGR3 | synaptogyrin 3 |
| 54985 | HCFC1R1 | host cell factor C1 regulator 1 |
| 89927 | C16orf45 | chromosome 16 open reading frame 45 |
| 5347 | PLK1 | polo like kinase 1 |
| 4502 | MT2A | metallothionein 2A |
| 4490 | MT1B | metallothionein 1B |
| 57555 | NLGN2 | neuroligin 2 |
| 1949 | EFNB3 | ephrin B3 |
| 247 | ALOX15B | arachidonate 15-lipoxygenase, type B |
| 5914 | RARA | retinoic acid receptor alpha |
| 4804 | NGFR | nerve growth factor receptor |
| 3675 | ITGA3 | integrin subunit alpha 3 |
| 332 | BIRC5 | baculoviral IAP repeat containing 5 |
| 125476 | INO80C | INO80 complex subunit C |
| 1058 | CENPA | centromere protein A |
| 9938 | ARHGAP25 | Rho GTPase activating protein 25 |
| 51263 | MRPL30 | mitochondrial ribosomal protein L30 |
| 90557 | CCDC74A | coiled-coil domain containing 74A |
| 6335 | SCN9A | sodium voltage-gated channel alpha subunit 9 |
| 3772 | KCNJ15 | potassium voltage-gated channel subfamily J member 15 |
| 29114 | TAGLN3 | transgelin 3 |
| 5010 | CLDN11 | claudin 11 |

| | | |
|---|---|---|
| 9982 | FGFBP1 | fibroblast growth factor binding protein 1 |
| 11162 | NUDT6 | nudix hydrolase 6 |
| 3148 | HMGB2 | high mobility group box 2 |
| 134147 | CMBL | carboxymethylenebutenolidase homolog |
| 81848 | SPRY4 | sprouty RTK signaling antagonist 4 |
| 8352 | HIST1H3C | histone cluster 1 H3 family member c |
| 3008 | HIST1H1E | histone cluster 1 H1 family member e |
| 11270 | NRM | nurim (nuclear envelope membrane protein) |
| 3159 | HMGA1 | high mobility group AT-hook 1 |
| 4188 | MDFI | MyoD family inhibitor |
| 7422 | VEGFA | vascular endothelial growth factor A |
| 6624 | FSCN1 | fascin actin-bundling protein 1 |
| 2115 | ETV1 | ETS variant 1 |
| 644150 | WIPF3 | WAS/WASL interacting protein family member 3 |
| 3638 | INSIG1 | insulin induced gene 1 |
| 286151 | FBXO43 | F-box protein 43 |
| 90865 | IL33 | interleukin 33 |
| 883 | KYAT1 | kynurenine aminotransferase 1 |
| 5355 | PLP2 | proteolipid protein 2 |

**Table S3:** Human Phospho-kinase array coordinates (ARY003B, RnDSystems).

| Membrane/ Coordinate | Target/Control | Phosphorylation Site |
|---|---|---|
| A-A1, A2 | Reference Spot | Ø |
| A-A3, A4 | p38α | T180/Y182 |
| A-A5, A6 | ERK1/2 | T202/Y204, T185/ Y187 |
| A-A7, A8 | JNK 1/2/3 | T183/Y185, T221/ Y223 |
| A-A9, A10 | GSK-3α/β | S21/S9 |
| B-A13, A14 | p53 | S392 |
| B-A17, A18 | Reference Spot | Ø |
| A-B3, B4 | EGF R | Y1086 |
| A-B5, B6 | MSK1/2 | S376/S360 |
| A-B7, B8 | AMPKα1 | T183 |
| A-B9, B10 | Akt 1/2/3 | S473 |
| B-B11, B12 | Akt 1/2/3 | T308 |
| B-B13, B14 | p53 | S46 |
| A-C1, C2 | TOR | S2448 |
| A-C3, C4 | CREB | S133 |
| A-C5, C6 | HSP27 | S78/S82 |
| A-C7, C8 | AMPKα2 | T172 |
| A-C9, C10 | β-Catenin | Ø |
| B-C11, C12 | p70 S6 Kinase | T389 |
| B-C13, C14 | p53 | S15 |
| B-C15, C16 | c-Jun | S63 |
| A-D1, D2 | Src | Y419 |
| A-D3, D4 | Lyn | Y397 |
| A-D5, D6 | Lck | Y394 |
| A-D7, D8 | STAT2 | Y689 |
| A-D9, D10 | STAT5a | Y694 |
| B-D11, D12 | p70 S6 Kinase | T421/S424 |
| B-D13, D14 | RSK1/2/3 | S380/S386/S377 |
| B-D15, D16 | eNOS | S1177 |
| A-E1, E2 | Fyn | Y420 |
| A-E3, E4 | Yes | Y426 |
| A-E5, E6 | Fgr | Y412 |
| A-E7, E8 | STAT6 | Y641 |
| A-E9, E10 | STAT5b | Y699 |
| B-E11, E12 | STAT3 | Y705 |
| B-E13, E14 | p27 | T198 |
| B-E15, E16 | PLC-γ1 | Y783 |
| A-F1, F2 | Hck | Y411 |
| A-F3, F4 | Chk-2 | T68 |
| A-F5, F6 | FAK | Y397 |
| A-F7, F8 | PDGF Rβ | Y751 |
| A-F9, F10 | STAT5a/b | Y694/Y699 |
| B-F11, F12 | STAT3 | S727 |
| B-F13, F14 | WNK1 | T60 |
| B-F15, F16 | PYK2 | Y402 |
| A-G1, G2 | Reference Spot | Ø |
| A-G3, G4 | PRAS40 | T246 |
| A-G9, G10 | PBS (Negative Control) | Ø |
| B-G11, G12 | HSP60 | Ø |
| B-G17, G18 | PBS (Negative Control) | Ø |

**<u>Table S4:</u>** Sequence of the C-terminal region of GPRC5A. Sequence was then cloned into a pET30a overexpressing plasmid, associated with some poly(Lysine/Arginine) tag (polyRK, light blue) to addressed it to the nucleus, a 6 Histidines tag (6His, green) to facilitate the purification of the polypeptide and separated by a sequence recognized by the TEV protease (TEV, grey).

| | |
|---|---|
| GPRC5A C-terminal DNA sequence | $^{5'}$acaaagcaacgaaaccccatggattatcctgttgaggatgctttctgtaaacctcaactcgtgaagaagagctatggt<br>gtggagaacagagcctactctcaagaggaaatcactcaaggttttgaagagacaggggacacgctctatgcccccta<br>ttccacacattttcagctgcagaaccagcctccccaaaaggaattctccatcccacgggcccacgcttggccgagccc<br>ttacaaagactatgaagtaaagaaagagggcagctaa$^{3'}$ |
| Translated coding sequence | $^{NH2}$TKQRNPMDYPVEDAFCKPQLVKKSYGVENRAYSQEEITQGFEETGDTL<br>YAPYSTHFQLQNQPPQKEFSIPRAHAWPSPYKDYEVKKEGS*$^{COOH}$ |
| Mimetic peptide sequence | MTKQRNPMDYPVEDAFCKPQLVKKSYGVENRAYSQEEITQGFEETGDTL<br>YAPYSTHFQLQNQPPQKEFSIPRAHAWPSPYKDYEVKKEGSRKRKRKRKR<br>KRKVDENLYFQGRGGGHHHHHHAAALEHHHHHH |

**<u>Table S5</u>**: Sequence of the plasmid surexpressing the C-terminal region of GPRC5A linked to GFP tag. GPRC5A C-ter sequence (blue) associated with the polyRK tag (light blue) was amplified from the previous construction (Table S3) and cloned into a pEGFP-C1 overexpressing plasmid, downstream of the coding sequence of GFP (green fluorescent protein, in green).

| | |
|---|---|
| pEGFP-GPRC5A-Cter DNA sequence | atggtgagcaagggcgaggagctgttcaccggggtggtgcccatcctggtcgagctggacggcgacgtaaacggccac<br>aagttcagcgtgtccggcgagggcgagggcgatgccacctacggcaagctgaccctgaagttcatctgcaccaccggc<br>aagctgcccgtgccctggcccaccctcgtgaccaccctgacctacggcgtgcagtgcttcagccgctaccccgaccacat<br>gaagcagcacgacttcttcaagtccgccatgcccgaaggctacgtccaggagcgcaccatcttcttcaaggacgacggc<br>aactacaagacccgcgccgaggtgaagttcgagggcgacaccctggtgaaccgcatcgagctgaagggcatcgacttc<br>aaggaggacggcaacatcctggggcacaagctggagtacaactacaacagccacaacgtctatatcatggccgacaag<br>cagaagaacggcatcaaggtgaacttcaagatccgccacaacatcgaggacggcagcgtgcagctcgccgaccactac<br>cagcagaacacccccatcggcgacggccccgtgctgctgcccgacaaccactacctgagcacccagtccgccctgagc<br>aaagaccccaacgagaagcgcgatcacatggtcctgctggagttcgtgaccgccgccgggatcactctcggcatggacg<br>agctgtacaagtccggactcagatctcgacggtaccgcgggcccgatatacatatgacaaagcaacgaaaccccatggat<br>tatcctgttgaggatgctttctgtaaacctcaactcgtgaagaagagctatggtgtggagaacagagcctactctcaagagg<br>aaatcactcaaggttttgaagagacaggggacacgctctatgcccccctattccacacattttcagctgcagaaccagcctcc<br>ccaaaaggaattctccatcccacgggcccacgcttggccgagcccttacaaagactatgaagtaaagaaagagggcagc<br>cgtaaacgtaagcgtaaacgtaaacgtaaacgtaaa |
| Translated coding sequence | MVSKGEELFTGVVPILVELDGDVNGHKFSVSGEGEGDATYGKLTLKFICTTG<br>KLPVPWPTLVTTLTYGVQCFSRYPDHMKQHDFFKSAMPEGYVQERTIFFKDD<br>GNYKTRAEVKFEGDTLVNRIELKGIDFKEDGNILGHKLEYNYNSHNVYIMAD<br>KQKNGIKVNFKIRHNIEDGSVQLADHYQQNTPIGDGPVLLPDNHYLSTQSAL<br>SKDPNEKRDHMVLLEFVTAAGITLGMDELYKSGLRSRRYRGPDIHMTKQRNP<br>MDYPVEDAFCKPQLVKKSYGVENRAYSQEEITQGFEETGDTLYAPYSTHFQL<br>QNQPPQKEFSIPRAHAWPSPYKDYEVKKEGSRKRKRKRKRKRKPGIHRI |
